# Fast Myosin Binding Protein-C is a Vital Regulator in Young and Aged Fast Skeletal Muscle Homeostasis

**DOI:** 10.1101/2024.12.26.629832

**Authors:** Akhil Baby, Kalyani Ananthamohan, Taejeong Song

## Abstract

**Background:** Skeletal muscle plays a vital role in voluntary motion and locomotion. Fast-twitch muscle fibers are characterized by their rapid contraction kinetics, high force generation capacity, and a distinct gene expression profile compared to slow-twitch fibers. Skeletal myosin binding protein-C (MyBP-C) paralogs, slow (sMyBP-C) and fast (fMyBP-C), interact with myosin and actin filaments within sarcomeres to modulate force development during contraction. These paralogs are differentially expressed in muscle fibers, with fMyBP-C predominantly expressed in the fast-twitch fibers. However, the role of fMyBP-C in skeletal muscle disease states and aging remains poorly understood. This study employs mouse models with fMyBP-C ablation to investigate its significance in skeletal muscle physiology.

**Methods:** Adult skeletal muscle samples aged 2∼7 months from male and female wild-type, db/db, MDX, ECC injury model, were used to determine the differential expression of fMyBP-C. Next, *Mybpc2* knockout (C2^-/-^) young (3∼5 months) and old (22 months) male mice were used to define the role of fMyBP-C in aging. Western immunoblotting was employed to analyze the expression of fMyBP-C and sMyBP-C and the phosphorylation status of sMyBP-C. The impact of C2^-/-^ and aging on the fiber type, size, and number as well as general muscle structure was assessed by immunohistochemistry and electron microscopy. The functional effect of C2^-/-^ and aging was measured in terms of in vivo and ex vivo muscle force generation. Lastly, RNA sequencing was performed to identify the molecular pathways dysregulated in the C2^-/-^ mediated muscle dysfunction in young and old mice.

**Results:** fMyBP-C was significantly reduced with a modest compensatory upregulation of sMyBP-C in the diseased fast-twitch muscles. fMyBP-C has a significantly higher expression in the male skeletal muscles compared to females. Further studies using young male C2^-/-^ mice showed a significant reduction in isometric tetanic force generation and relaxation rate, fiber type switching, atrophy, and altered gene expressions related to muscle function and metabolism compared to wild-type mice. Similarly, compared to their wild-type counterparts, aged male C2^-/-^ mice display significant deficits in muscle strength, endurance, and survival rate, accompanied by changes in muscle fiber size and molecular signaling pathways critical for muscle homeostasis.

**Conclusion:** fMyBP-C is an important regulator of muscle function and homeostasis in young and aged male fast-twitch muscle fibers. The absence of fMyBP-C aggravates the effect of aging on muscle structure and function. fMyBP-C has the potential to be a therapeutic target to modulate muscle wasting caused by aging and disease.

## Introduction

Fast-twitch muscle fibers, characterized by their rapid contraction kinetics and high force generation capacity, are indispensable for executing rapid and forceful body movements. Compared to the slow-twitch, fast-twitch fibers have different compositions of key sarcomere elements such as myosin isoform (heavy and light chains) and troponin complex. Myosin binding protein-C (MyBP-C) is a sarcomeric regulatory protein located in the C-zone of the sarcomere’s I-band. It interacts with myosin and actin filaments within sarcomeres, modulating cross-bridge cycling kinetics and regulating force development during muscle contraction [1, 2]. MyBP-C is also one such protein having a distinct expression profile in slow- and fast-twitch muscles [3].

Among the three MyBP-C paralogs, fast MyBP-C (fMyBP-C) expression is predominant in fast-twitch muscle fibers, particularly in type IIb, with some presence in type IIx fibers. This protein plays a substantial role in enhancing both muscle contractility and structural integrity [4, 5]. This fine-tuning of contractile machinery in fast-twitch fibers is crucial for precisely regulating muscle force and speed. Dysregulation or loss of fast MyBP-C disrupts this control, leading to impaired muscle function, decreased force production, and a potential decline in motor performance [4]. Consequently, understanding the role of fast MyBP-C is essential for elucidating mechanisms underlying muscle contractility and its impact on overall muscle function. fMyBP-C also contributes to sarcomere stability and maintaining their ordered arrangement. Disruptions in fMyBP-C expression can lead to structural abnormalities, including myofilament misalignment and disorganization. Such structural alterations compromise muscle integrity, rendering the fibers susceptible to injury and functional impairment [4, 6]. Therefore, the preservation of fMyBP-C is crucial for maintaining the structural integrity of fast-twitch muscles.

During aging, disruptions in the fast-twitch muscle homeostasis can significantly impact both muscle function and overall health. Aging is invariably associated with a decline in muscle mass and function, a condition known as sarcopenia. Notably, fast-twitch muscle fibers are particularly susceptible to age-related changes, including fiber atrophy and a reduction in contractile functions [7, 8]. However, the precise role of fMyBP-C in maintaining homeostasis within aging fast-twitch muscles remains unknown. It is conceivable that the loss of fMyBP-C and the resultant alterations in muscle structure and function may be instrumental in diminishing muscle force generation and increasing muscle damage, thereby amplifying the morbidity experienced by older individuals.

In this study, we have conducted a comprehensive profiling of fMyBP-C expression across various disease conditions and assessed the impact of fMyBP-C deletion on muscle homeostasis and age-related muscle loss. Our investigation reveals notable variations in fMyBP-C expression within different muscle conditions and among male and female mice. Additionally, our findings highlight the crucial role of fMyBP-C in maintaining muscle homeostasis, and we have established its essential contribution to preserving muscle function and structural integrity in aging muscles.

## Materials and Methods

### *Mybpc2* knockout *(Mybpc2* KO) mouse model

Two *Mybpc2* knockout (KO) mouse models were used in this study. The first model (FVBN background) was generated by the targeted replacement of *Mybpc2* exon 2 to 22 with a Neo cassette flanked by two LoxP sites, as previously documented (*1*). The second *Mybpc2* KO mouse model (C57BL/6 background) was created by applying the CRISPR/Cas9 technology, targeting exons 6 and 7 of the *Mybpc2* gene. This approach resulted in indel mutations and the deletion of *Mybpc2*, as illustrated in *Figure S1*. The second mouse model was used for in vivo plantar flexor function tests in young mice and for PKA-dependent sMyBP-C phosphorylation experiments. All other data were collected from *Mybpc2* KO mice with an FVBN background. The diabetic (db/db) and Duchenne muscular dystrophy (MDX) mice (3∼7 months old) utilized in this study were purchased from Jackson Laboratory. For all experimental procedures, mice were anesthetized using 2.0-2.5% isoflurane inhalation and euthanized by cervical dislocation before the tissue collections. All animal-related protocols were conducted in strict compliance with the guidelines approved by the Institutional Animal Care and Use Committee at the University of Cincinnati.

### Muscle strength tests

#### Grip strength

Forelimb grip strength was assessed through three trials, during which the mouse was gently pulled against a grip strength meter (1027SM, Columbus Instruments). The maximum recorded value was normalized to the respective body weight and subjected to group-wise comparisons.

#### In vivo plantar flexor force generation

The isometric tetanic torque of the plantar flexor muscle was assessed in vivo, following the previously established protocol [9]. With the mouse under anesthesia using 2.0% inhaled isoflurane, the right knee was immobilized at a 90° angle, and the foot was securely fastened to a footplate connected to a dual-mode servomotor (Model 300C-LR, Aurora Scientific). Following the determination of the peak isometric twitch force generation, the isometric tetanic force was recorded at electrical frequencies ranging from 25 to 150 Hz for 350 milliseconds at intervals of two minutes using an Aurora Scientific apparatus (Model 701C).

#### Ex vivo EDL muscle function

As previously outlined [4], ex vivo EDL muscle contractile properties were assessed in a tissue bath containing oxygenated Krebs-Henseleit buffer (pH 7.4) at 36°C. The proximal tendon was fastened to a secure pin, while the distal tendon was connected to a dual-mode lever system using silk sutures. Before electrical stimulation, the muscle was allowed to equilibrate in a relaxed state within the bath for 10 minutes. After determining the peak isometric twitch force (Pt), we proceeded to measure the peak isometric tetanic force (Po) through a series of stimulations ranging from 25 to 200 Hz, with intervals of two minutes.

A fatigue test was performed by subjecting the muscle to repeated contractions at 150 Hz, with each contraction separated by 10-second intervals, a total amounting to 10 contractions. The physiological cross-sectional area (CSA) of the muscle was calculated by dividing muscle weight (in grams) by the product of fiber density (1.06 mg/mm^3^) and optimal fiber length (in millimeters). The optimal fiber length was determined by dividing the muscle length by a ratio of 0.44, as described previously [10]. Specific force (SPo) was computed by normalizing Po by CSA (Po/CSA) and expressed in units of N/cm^2^. Muscle function data, both in vivo and ex vivo, were digitized using Dynamic Muscle Control (DMC v5.5) software and subsequently analyzed employing Dynamic Muscle Analysis (DMA v5.3) software from Aurora Scientific.

### Eccentric contraction (ECC) induced muscle injury

To induce muscle injury, we administered repeated eccentric muscle contractions (ECC, n=100) in the right hindlimb by forcibly inducing dorsiflexion while concurrently generating maximum isometric tetanic torque in the plantar flexors at a frequency of 150 Hz for a duration of 350 milliseconds, with a 5-second interval between contractions. This was executed at an angle of 14 degrees, with a speed of 0.7 degrees per 10 milliseconds as previously described [4]. Muscle samples were collected from the gastrocnemius, soleus, and plantaris muscles of the injured leg at time points of 0.5, 1.0, 3.0, and 48 hours post-injury. Contralateral uninjured muscle samples collected at 0.5 hours after the injury were served as a control.

### Western blot analyses

The level of total and phosphorylated proteins was determined following established procedures [4]. Briefly, 10-20 μg of homogenized protein samples were loaded onto pre-casting gels (Bio-Rad) and electrophoresed at 80-100V for 90 minutes. Subsequently, the samples were transferred onto nitrocellulose membranes and blocked in 5% non-fat dry milk for 1 hour. Primary antibodies, including sMyBP-C (WH0004604M1 and SAB3501005, Sigma-Aldrich), fMyBP-C (SAB2108180, Sigma-Aldrich), Myh4 (BF-F3, DSHB), MyoM1 (20360-1-AP, Proteintech), Mylpf (A24975, Antibodies.com), Foxo1 (18592-1-AP, Proteintech), Ankrd2 (11821-1-AP, Proteintech), Cryab (MA5-27708, Invitrogen), and phospho-sMyBP-C (S59/S62 and T190/T191, ProSci Inc, Order # PAS22457 and #PAS22459), were incubated overnight at 4°C. Phospho-specific sMyBP-C antibodies targeting Serine 59, Serine 62, and Serine 204 were generously provided by Dr. Kontrogianni-Konstantopoulos (University of Maryland School of Medicine). Secondary antibodies conjugated with infrared fluorescent dye were incubated for 1 hour at room temperature, and protein expression was detected and quantified using the Odyssey CLx system (Li-Cor). β-actin (3779, ProSci) served as a loading control.

### ELISA

Standards for sMyBP-C (BIOMATIK, Cat# EKU06086) and fMyBP-C ELISA (abbexa, Cat# abx534308) provided in the ELISA kits were resuspended in Standard Diluent buffer to create a 10 ng/ml Standard Solution. Serial dilutions of this stock were prepared to yield concentrations of 0.078, 0.156, 0.312, 0.625, 1.25, 5, and 10 ng/mL. Samples were diluted in PBS to 100µL to standardize volumes, and the dilution factor for final calculations was determined. Both samples and standards were brought to room temperature before use. Duplicates of standards, test samples, and control (zero) wells were set on antibody pre-coated plates, and their positions were noted. Solutions were added to each well bottom without touching the sides. Standards and samples were gently mixed before addition, avoiding foaming. 100 µl of diluted standards were added to standard wells, 100 µl of Standard Diluent buffer to control wells, and 100 µl of appropriately diluted samples to test sample wells. Plates were mixed, sealed, and incubated for 2 hrs at 37°C. After discarding liquids, 100 µl of Detection Reagent A was added to each well and incubated for 1 hr at 37°C. Plates were washed thrice with 1X Wash Buffer, removing any remaining buffer. 100 µl of Detection Reagent B was added to each well and incubated for 1 hr at 37°C. The wash process was repeated five times, followed by the addition of 90 µl of TMB Substrate to each well and incubating at 37°C for 15 minutes. 50 µl of the Stop Solution was added to each well and mixed gently. Thereafter, the absorbance (OD) was measured at 450 nm. Relative OD was calculated as (OD of Each Well) – (OD of Zero Well). A standard curve was plotted using relative OD against the respective concentration of each standard. Sample concentrations were interpolated from the standard curve, and to factor in sample dilution, the concentrations were adjusted by the dilution factor.

### PKA-dependent sMyBP-C phosphorylation

Dissected EDL muscles from WT and C2^-/-^ mice were skinned in a glycerinated relaxing solution containing the following components (in mM): 7 EGTA, 100 BES, 0.017 CaCl2, 5.49 MgCl2, 5 DTT, 15 creatine phosphate, 4.66 ATP, 55.7 K-propionate; pH 7.0, and 50% (v/v) glycerol. This was performed at 4°C on a gently rotating shaker for 24 hours. Muscle tendons were affixed to a wooden stick using silk sutures to secure resting muscle length. The glycerinated relaxing solution was changed thrice during the 24 hours, and skinned samples were subsequently stored at -20°C. Four heads of the EDL were then separated and cut into small bundles along their long axis in a relaxing solution. After being washed three times with PBS, the samples were incubated in 200uL of PKA (0.5U/uL, Sigma-Aldrich, P2645) for 1 hour at room temperature on a shaker. Skinned muscle fibers were homogenized, and sMyBP-C phosphorylation was probed using specific antibodies following the aforementioned protocol.

### Immunohistochemistry

Immediately after dissection, the EDL muscles were embedded in O.C.T. compound and flash-frozen in isopentane cooled with liquid nitrogen. Cross-sectioned slides, with a thickness of 10 μm, were prepared and stained with H&E or antibodies targeting myosin heavy chains (MHC type I, BA-D5; MHC type IIA, SC-71; MHC type IIb, BF-F3 from DSHB) and laminin (L9393 from Sigma-Aldrich). Secondary antibodies conjugated with Alexa Fluor (Invitrogen) were used for immunohistology. All images were acquired using a Leica DMi8 microscope and analyzed with the ImageJ (NIH) software in a blinded manner.

### Transmission electron microscopy

EDL muscles were fixed through whole-body perfusion with a solution of 3% paraformaldehyde and 0.1% glutaraldehyde in a 0.1M cacodylate buffer for 2 hrs. Following dissection, the muscles were submerged for 2 hrs in a solution of 2.5% glutaraldehyde in a 0.1 M cacodylate buffer with a pH of 7.4 at room temperature. Subsequently, the fixed muscles were sectioned into smaller pieces and subjected to post-fixation using 1% w/v OsO4 dissolved in distilled water. The muscles were then dehydrated gradually using a series of ethanol solutions and finally embedded in Epon resin. 70 nm thickness sections were prepared and initially stained with uranyl acetate followed by lead citrate. Specimens were examined and imaged using a Tecnai G2 Spirit transmission electron microscope.

### RNA sequencing analysis

The quantification of raw gene counts was carried out through the utilization of feature Counts (v1.5.2) (15), and subsequent normalization was achieved by applying edgeR’s TMM (trimmed mean of M values) method. To identify differentially expressed genes (DEGs), we employed the limma/voom approach. Specifically, genes surpassing the defined moderate threshold criteria (|fold| > 1.5X and (adjusted) p < 0.05) were designated as differentially expressed. These DEGs were then employed as queries in EnrichR and ClueGO to retrieve enriched pathways and/or gene ontology terms. Additionally, we performed a GSEA analysis using gene-set permutation. Additional functional enrichment analysis for DEGs specific and common to young and old C2-/-mice has been carried out using metascape [11]. The DEGs and their enrichment data are available in the supplementary file, SuppInfo.xls.

### Statistics

The data is reported as the mean ± standard error (SE). Comparisons were conducted using either Student’s t-test for experiments with two groups or one-way ANOVA with Tukey’s post-hoc test for more than two groups, utilizing GraphPad Prism 7.04 software. Survival curves of WT and C2^-/-^ mice were assessed with the Mantel-Cox test. Statistical significance was defined as a P-value below 0.05.

## Results

### Reduced fMyBP-C protein expression in diseased and injured muscles

fMyBP-C expressed in the sarcomeres of fast-twitch muscle fibers is required for modulating the kinetics of the cross-bridge cycle and maintaining sarcomere integrity, allowing the precise control of muscle force and speed during rapid high-force generation [4, 6]. In this study, we first evaluated whether the protein expression of the two skeletal MyBP-C paralogs (Figure 1A), sMyBP-C and fMyBP-C were altered in diseased and injured muscles. We found a significant reduction of fMyBP-C in the tibialis anterior (TA) muscle of db/db mice. However, sMyBP-C levels were significantly elevated in the TA muscle of both db/db and MDX mice models (Figure 1B-D). As a result, the ratio of sMyBP-C to fMyBP-C was significantly increased in both diseased muscles compared to the wild-type (WT) controls (Figure 1E). In gastrocnemius muscle exposed to eccentric contraction-induced (ECC) muscle injury, we observed a notable reduction of fMyBP-C protein level at time-points, 3 and 48 hrs post-injury. Whereas sMyBP-C protein exhibited a more modest decline following the injury (Figure S2C). Notably, fMyBP-C was released to circulation and detected in serum samples at 1 and 24 hrs after ECC injury, but we did not see a detectable amount of sMyBP-C in the same samples (Figure S2D). However, when the muscles at different time points post-injury were incubated in PBS, fMyBP-C and to a lesser extent sMyBP-C were detected in the 6.0-hour effluent sample (Figure S2E and F). These findings collectively indicate that both absolute and relative fMyBP-C expression levels were compromised in diseased and injured muscle tissues, and loss of fMyBP-C may contribute to the functional deficits in these muscles.

**Figure 1.**
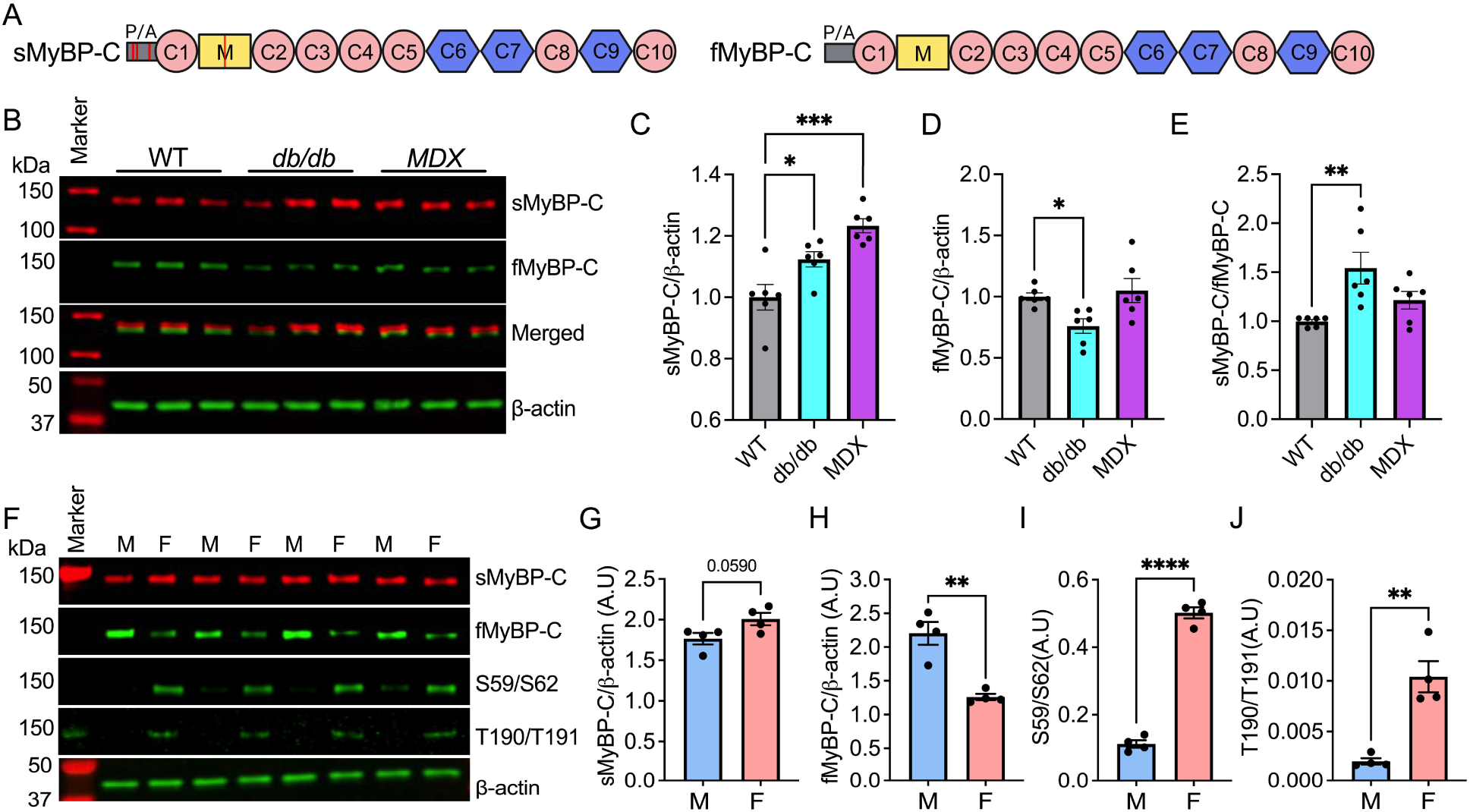
Reduced fMyBP-C protein level in diseased muscles and differential fMyBP-C expression in male and female skeletal muscle. (A) sMyBP-C and fMyBP-C domain structure (P/A: proline and alanine-rich region, red circle: immunoglobulin-like domain, blue hexagon: fibronectin type3 domain, red line: phosphorylation site). (B) Altered sMyBP-C and fMyBP-C protein expressions in wild-type control (WT) versus diseased (db/db and MDX) tibialis anterior muscles measured by Western blot analysis. Quantitative comparison of the protein expressions between groups (C-E). n = 6 mice TA muscles. WT vs. db/db and MDX. Represent Western blot images (F), quantifications of sMyBP-C (G), fMyBP-C (H), and phosphorylation of sMyBP-C at Serine 59/62 and Threonine 190/191 (I-J) in male and female mouse plantaris muscles (n = 4 in each). M denotes male and F denotes female group. Error bars represent ± SEM and \**P*<0.05, \*\**P*<0.01, and \*\*\*\**P*<0.0001.

### Distinct slow and fast MyBP-C expression profile in male and female muscles

Our previous study demonstrated the distinctive expression of fMyBP-C in fast-twitch muscle fibers, and it is well-established that these fibers exhibit higher expression levels in male skeletal muscle compared to female skeletal muscle [4, 12]. Therefore, we have conducted an assessment of slow and fast MyBP-C protein expressions, as well as the phosphorylation status of sMyBP-C, in male and female muscle samples. Our findings revealed a notable disparity, with male plantaris muscle exhibiting a significantly higher (75% more) fMyBP-C expression when compared to their female counterparts (Figure 1F-H). Conversely, sMyBP-C phosphorylation at Ser59/Ser62 and Thr190/Thr191 were significantly higher in the female samples, with increases of 4.42 and 5.08 fold, respectively, compared to the male samples (Figure 1I and J). These results underscore the significance of fMyBP-C in male and female muscle physiology and, highlight its greater importance in the function and structural integrity of male fast-twitch fibers. Consequently, our subsequent experiments primarily investigate the consequences of fMyBP-C deletion in the male skeletal muscle.

### Reduced force generation capacity and disrupted muscle fiber architecture in the absence of fMyBP-C

To examine the role of fMyBP-C in the contractile function of skeletal muscle, we measured the in vivo isometric plantar flexor force generation in male WT and C2^-/-^ mice. As shown in the force-frequency graph (Figure 2A), there was no difference in force generation between WT and C2^-/-^ when generating incomplete tetanic contraction at low electrical frequencies (25 and 50Hz). However, at a frequency of 75 Hz and above, which induces tetanic muscle contraction, C2^-/-^ generated significantly less force than WT. Peak twitch force generation was equivalent in both groups, but peak tetanic force was significantly lower (25% less) in C2^-/-^ compared to WT (Figure 2B and C). Half relaxation time and rate of activating during peak tetanic contraction were not different between groups, but the rate of relaxation was significantly slower (42% less) in C2^-/-^ (Figure 2D and E).

**Figure 2.**
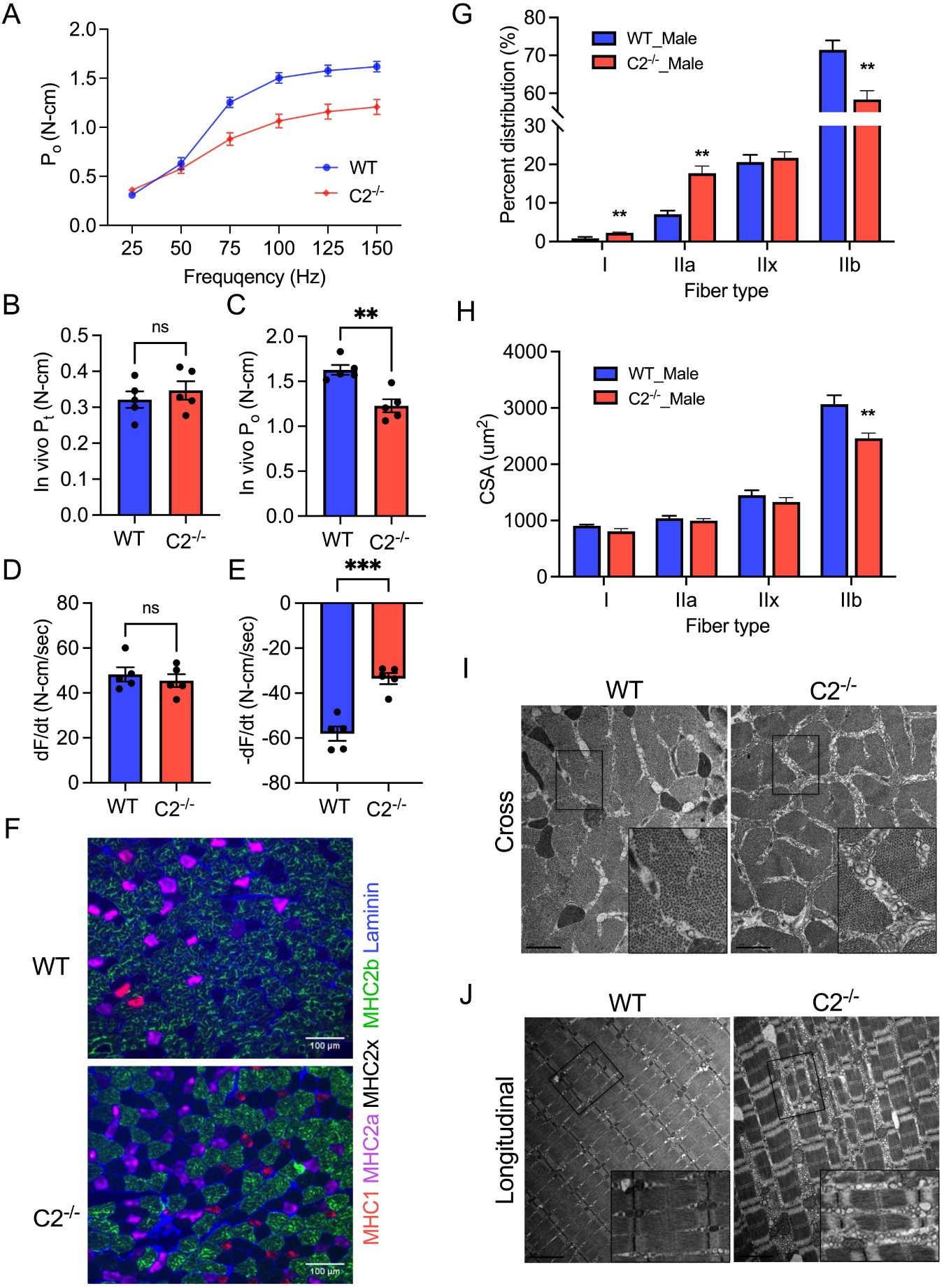
Impaired muscle function and structure in male C2^-/-^ muscle. A. In vivo isometric plantar flexor force generation of male WT and C2^-/-^ mice at different electrical frequencies (25 to 150Hz). Peak twitch force (B), peak tetanic force (C), and the rate of activation/relaxation (D and E). (F) Cross-sectioned EDL muscles immuno-stained with MHC and laminin antibodies. The scale bar is 100µm. Fast to slow fiber type switch (G) and significant atrophy of type IIb fibers (H) in C2^-/-^ vs. WT muscle. n = 6 slides from three muscles in each group. Cross (I) and longitudinal (J) sectioned EM images of EDL muscles show the accumulation of small vesicles between myofibrils of C2^-/-^. The scale bar is 1µm (Cross image) and 2µm (Longitudinal image). Error bars indicate SEM. \*\**P*<0.01 and \*\*\**P*<0.001.

Histo-pathological adaptation of each muscle fiber type in the absence of fMyBP-C was evaluated in cross-sectioned EDL muscle stained with myosin heavy chain (MHC) antibodies (Figure 2F). Our results showed significant muscle fiber type switching from fast to slow fiber types in C2^-/-^. The percentage of type IIb fibers significantly decreased (12.8% less) while increasing the percentages of type I and IIa fibers, 1.4 and 10.6% respectively (Figure 2G). The average cross-sectional area (CSA) of each fiber type was also measured and a significant reduction in CSA of type IIb fibers (-20%) was observed in C2^-/-^ (Figure 2H). These results indicate that fMyBP-C is required to maintain the number and size of fast-twitch fiber (type IIb) in male skeletal muscle. Electron microscopy analysis of male EDL muscle presented preserved overall sarcomere structure and integrity in C2^-/-^ which is consistent with what was previously shown [4]. However, we found expanded space between muscle fibers and small vesicles occupied the expanded space possibly indicating ongoing muscle atrophy or chronic stress in the C2^-/-^ muscle (Figures 2I and J).

### Reduced PKA-dependent sMyBP-C phosphorylation in C2^-/-^ muscle

PKA-dependent MyBP-C phosphorylation is crucial in regulating muscle contraction, impacting muscle contractility and relaxation, thereby influencing overall muscle function [13]. sMyBP-C has multiple PKA-dependent phosphorylation sites in its N’-terminal, and our previous study showed a compensatory increase of sMyBP-C protein expression in C2^-/-^ EDL [4, 14]. Therefore, we have evaluated changes in sMyBP-C phosphorylation in skinned EDL fiber after PKA treatment. In wild-type (WT) samples, phosphorylation of sMyBP-C at serine 59 and serine 62 was significantly increased in PKA-treated fibers while no changes were observed for serine 204 phosphorylation (Figure 3A-D). However, in C2^-/-^ mice fibers, serine 59 and serine 62 sites were not responsive to PKA treatment. Instead, phosphorylation of the serine 204 site was significantly increased after PKA treatment (Figure 3F-I). As expected, the total sMyBP-C expression level was not different between PKA-treated and non-treated samples of WT and C2^-/-^ (Figure 3E and J).

**Figure 3.**
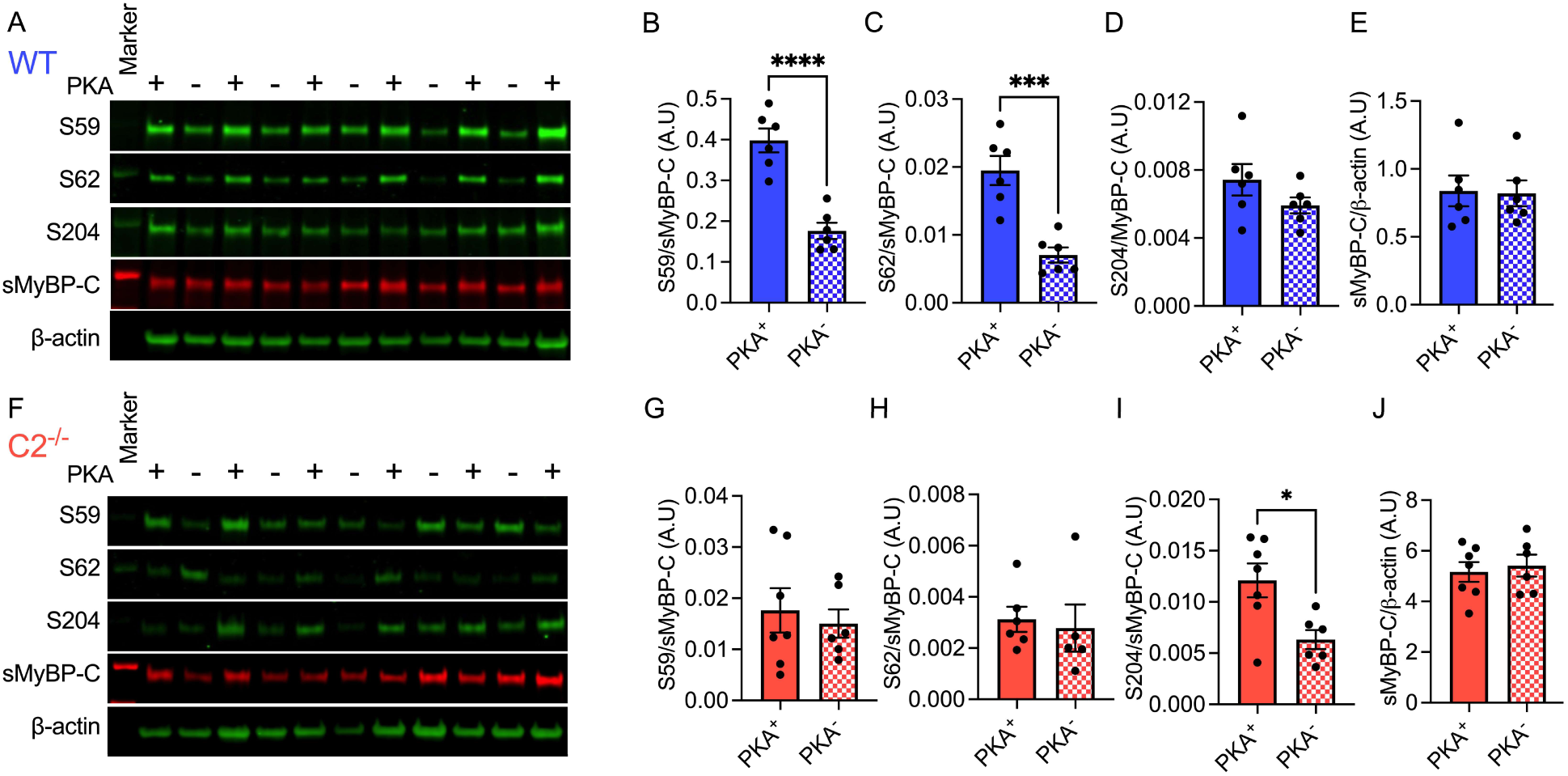
Altered PKA-dependent sMyBP-C phosphorylation in C2^-/-^ fiber. (A) PKA-targeted sMyBP-C phosphorylation was measured by Western blot in skinned WT (A) and C2^-/-^ (F) EDL fibers. Quantification of sMyBP-C phosphorylations at serine 59 (B and G), serine 62 (C and H), serine 204 (D and I), and total sMyBP-C (E and J) in WT and C2^-/-^ fibers. Error bars represent ± SEM and \**P*<0.05, \*\*\**P*<0.001, and \*\*\**P*<0.0001. A total of 6-7 skinned fiber preparations were used.

### Dysregulation of Critical genes and pathways in C2^-/-^ muscle

To further investigate the underlying molecular mechanisms of reduced contractile functions and structural integrity in male C2^-/-^ muscle, we have reanalyzed male single EDL fiber RNA Seq data (n = 10 in each group, GEO accession: GSE 160827). We found a total of 956 DEGs (621 up and 335 down) in C2^-/-^ (Figure 4A). The ten most up- and downregulated genes in C2^-/-^ fibers are listed in Figure 4B. The most upregulated genes included Actc1; a cardiac and developing skeletal muscle sarcomeric gene and Phlda3; a positive apoptotic pathway regulator. Membrane ion and protein transporter genes; Tm87a and Atp13a2 and Wnt signaling inhibitor gene; Apcdd1 are among the most downregulated genes in C2^-/-^ (Figure 4C). We also found significantly disrupted biological process (GO_BP) and molecular function (GO_MF) pathways using gene ontology analyses. Metabolic processes and membrane receptor binding pathways are up-regulated while down-regulating DNA and RNA regulation pathways (Figure 4D and E). Altogether, findings from C2^-/-^ mice collectively underscore fMyBP-C’s pivotal function in sustaining the structural integrity and contractile functionality of fast-twitch muscle fibers in male skeletal muscle.

**Figure 4.**
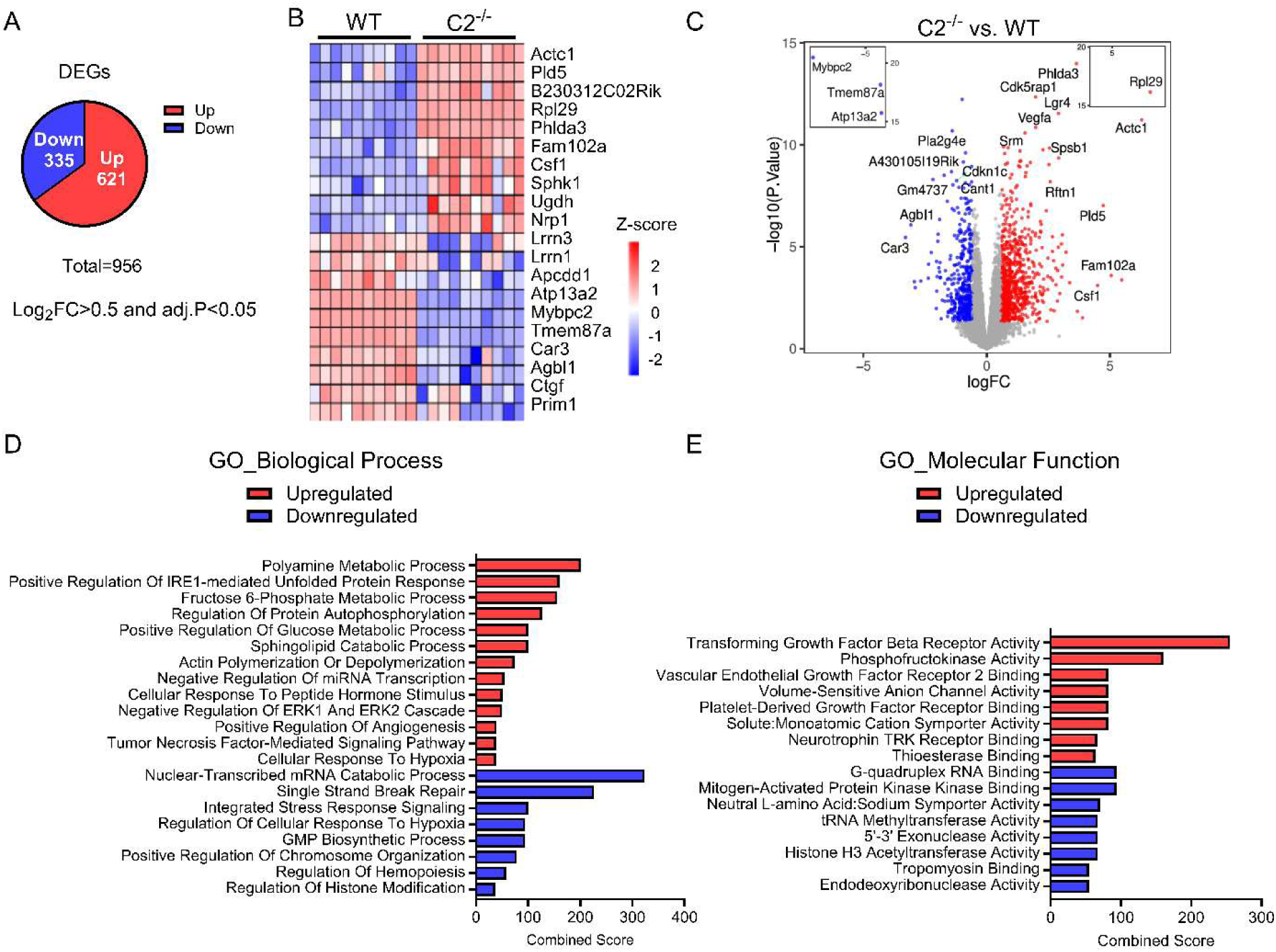
RNA sequencing reveals dysregulated genes and pathways in young male C2^-/-^ EDL muscle fiber. RNA Sequencing was carried out on EDL muscle samples from young male C2^-/-^ and wildtype (WT) mice (n = 10/group), followed by differential gene expression analysis and gene set enrichment analysis. (A) Total number of differentially expressed genes based on log-transformed fold change cut off of 0.5 and effect size threshold of adj.P<0.05. (B) Heatmap of top ten up and down-regulated genes, and (C) Volcano plot comparing DEGs in C2^-/-^ vs. WT. (D-E), Gene set enrichment analysis of DEGs revealed the top up and down-regulated biological processes (D) and molecular function (E) in C2^-/-^ vs. WT. EDL muscles.

### fMyBP-C is required for aged muscle function and homeostasis

Maintaining the integrity and functionality of skeletal muscle becomes increasingly challenging with age. Preserved muscle mass and strength have been known to decrease the risk of orthopedic injury and increase the quality of life and independence in the aged population [15]. Literature has shown selective loss of glycolytic type II fibers with aging. Particularly, type IIb fibers (2x in humans) are the most susceptible to atrophy, related to developing metabolic dysfunction in aged muscle [16–18]. Therefore, we investigated the roles of fMyBP-C in age-related loss of muscle functions and structures.

Gross body mass and muscle weight did not differ between WT^Old^ and C2^-/-Old^ mice (Figure S3). Although the survival rate was not significantly different between the two groups, C2^-/-Old^ mice showed a gradual decline with aging. At 20 months, 85% of WT mice were alive, while only 63% of C2^-/-Old^ mice survived (Figure 5A). In vivo, grip strength decreased with aging in both groups, but C2^-/-Old^ mice exhibited significantly lower grip strength, as well as in vivo peak isometric plantar flexor force at 22 months (Figures 5B and C). To further evaluate fast-twitch muscle functions, we isolated EDL muscle and measured ex vivo contractile functions which resulted in significantly reduced force generation capacity at the electrical stimulation frequency above 100Hz in C2^-/-Old^. Compared to WT^Old^ control, peak isometric tetanic and specific force significantly reduced by 21% and 26% in C2^-/-Old^, respectively. C2^-/-Old^ EDL also showed reduced fatigue resistance during repeated peak isometric tetanic contractions showing more Po loss after the fourth tetanic contraction (Figure 5D-G).

**Figure 5.**
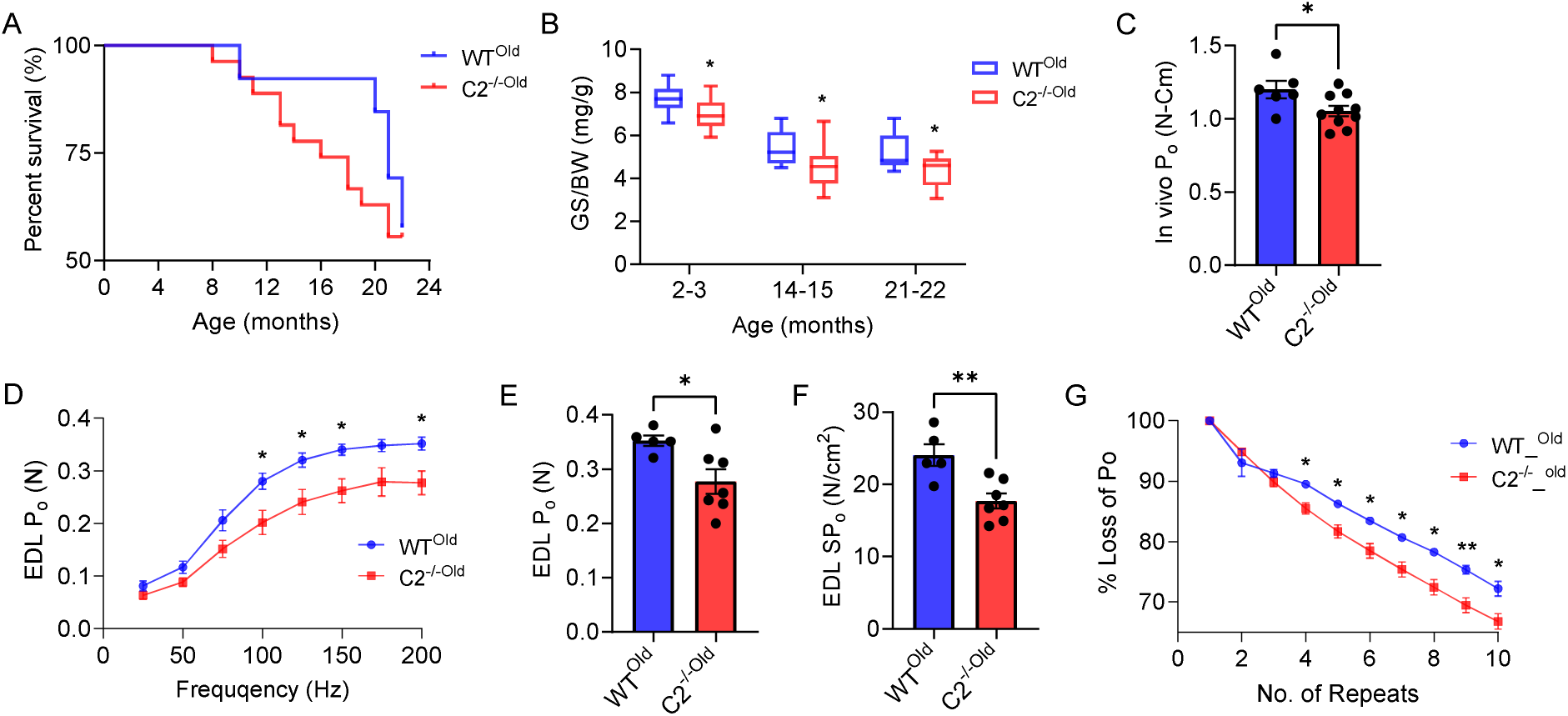
Reduced survival rate and muscle functions in C2^-/-^ mice. (A) Mouse survival rate decreased after 8 months of age in C2^-/-^ (n = 27) vs. WT (n = 13) mice. (B) Grip strength was significantly lower in C2^-/-^ (n = 15∼16) compared with WT (n = 7∼16) mice of all ages. (C) In vivo, peak isometric plantar flexor force generation was reduced in C2^-/-^ mice. (D) Downshift force-frequency graph. C2^-/-^ EDL generated significantly less isometric tetanic force at mid to high electrical frequency (100∼200Hz). Ex vivo EDL peak isometric tetanic (E) and specific force (F) were significantly lower in C2^-/-^ (vs. WT). (G) C2^-/-^ EDL loses significantly more force during repeated ten isometric peak tetanic contractions. n = 4∼10 in each group. Age = 2 to 22 months. Error bars represent ± SEM and \**P*<0.05 and \*\**P*<0.01 C2^-/-Old^ vs. WT^Old^.

Next, to evaluate histological adaptations in the aged WT and C2^-/-^ muscle, we first stained cross-sectioned EDL samples with H&E and counted the number of central nuclei (CN) which is a hallmark of muscle regeneration after damage (Figure 6A, top). The average numbers of CN in each slide were not different between the two groups, however, we found significantly increased fiber numbers in C2^-/-Old^ samples. The muscle fiber size distribution graph shifted left in C2^-/-Old^ indicating the increased number of small-sized muscle fibers (Figure 6B-D). EDL slides were also immunostained with MHC antibodies and analyzed the fiber type distribution as well as the size of each fiber type (Figure 6A, bottom). Muscle fiber type composition was not different in WT^Old^ and C2^-/-Old^ EDL muscles, but the size of fast-twitch fibers (type IIa, IIx, and IIb) was significantly reduced in C2^-/-Old^ (Figure 6E and F). These results emphasize that fMyBP-C ablation results in severe muscle pathology and that it is essential for maintaining fast-twitch fiber size and homeostasis during the aging process.

**Figure 6.**
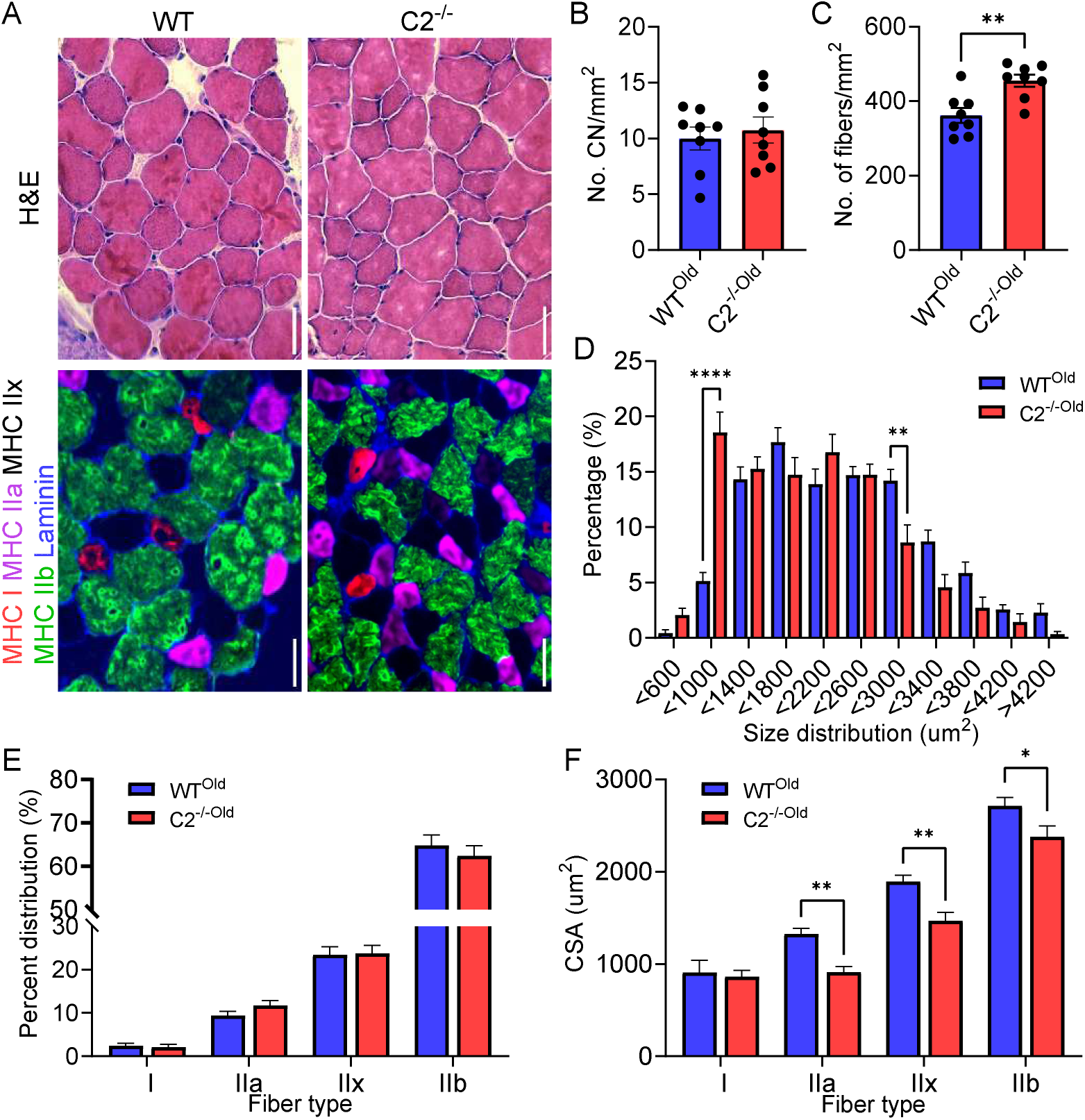
Atrophy of fast twitch fibers in aged C2^-/-^ EDL. (A) Represent H&E (top) and myosin heavy chain isoforms (MHC, bottom) images of WT (left) and C2^-/-^ EDL. Scale bar=50µm. There was no difference in the numbers of central nuclei (B) but a significant increase in the total number of fibers in C2^-/-^ compared to WT (C). (D) Increased percentage of small-sized fibers while decreasing large fibers in C2^-/-^. (E) Preserved fiber type distributions but (F) significantly reduced the cross-sectional area of fast twitch fibers (type IIa to IIb) in C2^-/-^. n = 6∼8 slides from 3-4 mice in each group. Error bars mean ± SEM and \**P*<0.05 and \*\**P*<0.01 C2^-/-Old^ vs. WT^Old^ fibers.

### Disrupted gene and protein expressions in aged C2^-/-^ muscle

To further explore the molecular mechanisms underlying the reduced muscle function and fiber size observed in aged C2^-/-^ muscle, we conducted an RNA expression profiling of the tibialis anterior (TA) muscle using RNA seq analysis. The top 20 most up- and down-regulated genes identified in aged C2^-/-^ fibers are listed in Figure 7A. Among the 310 DEGs (226 up and 84 down) in the aged samples (Figure 7B), the most dysregulated genes included Actc1 (up-regulated) and several genes with predicted but not yet validated for its functions, including Myh13 (up-regulated); myosin heavy chain 13, which is predicted to enable actin filament binding activity and microfilament motor activity, Fam174b (down-regulated), predicted to be involved in Golgi organization and expressed in hearts; and Apold1 (down-regulated), predicted to enable lipid binding activity and is expressed in cardiac ventricles (Figure 7C). Comparing the DEGs from C2^-/-^ and C2^-/-Old^ mice revealed common signatures (23 upregulated genes and 10 downregulated genes) specific to the absence of fMyBP-C (Figure S5A). Similarly, there is a distinct set of DEGs in C2^-/-^ and C2^-/-Old^ mice, implicating the role of fMyBP-C in the aging process. More importantly, gene set enrichment analysis (GSEA) revealed 328 upregulated and 104 downregulated pathways (Figure S5B) and illustrates the key pathways activated and suppressed (Figure S5C) and related genes (Figure S6 & S7), crucial for maintaining muscle function and homeostasis. Notably, pathways associated with collagen formation, ECM receptor interaction, and negative regulation of muscle differentiation, which are linked to muscle damage and impaired regeneration, were significantly increased in C2^-/-Old^ (Figure 7D-F). Concurrently, genes related to skeletal muscle contraction showed significant reductions in aged C2^-/-^ muscle (Figure 7G). We have also analyzed the expression of key proteins related to muscle structure and atrophy. Notably, the expression of sMyBP-C was significantly increased in aged C2^-/-^ muscles, similar to the levels observed in young C2^-/-^ mice. Interestingly, the expression of Myomesin-1 (Myom1), which plays a crucial role in stabilizing thick filaments at the M-band, also showed a significant increase in the aged C2^-/-^ samples. However, the protein levels of Myh4, Mylpf, Foxo1, Ankrd2, and Cryab did not exhibit any significant differences between the two groups (Figure S8).

**Figure 7.**
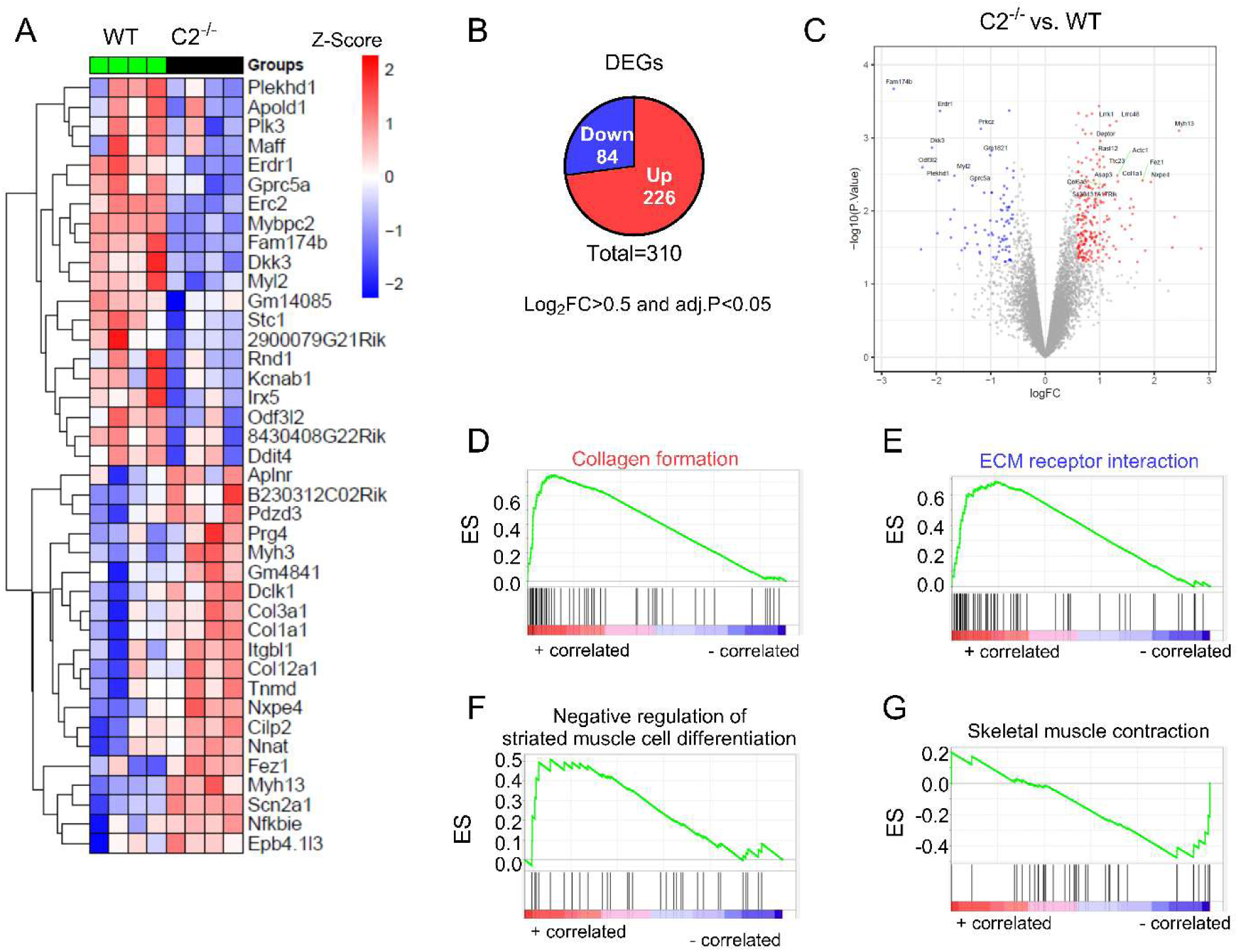
RNA sequencing reveals dysregulated genes and pathways in aged tibialis anterior C2^-/-^ TA muscle. RNA Sequencing was carried out on TA muscle samples from old male C2^-/-^ and wildtype (WT) mice (n = 4/group), followed by differential gene expression analysis and gene set enrichment analysis. (A) Heatmap of top twenty up and down-regulated genes, (B) Total number of differentially expressed genes based on log-transformed fold change cut off of 0.5 and effect size threshold of adj.P<0.05. (C) Volcano plot comparing DEGs in C2^-/-^ vs. WT. (D-G), Enrichment score graph of key molecular signatures that alters muscle structure and function in older male C2^-/-^ vs. WT TA muscles.

## Discussion

The sarcomere is a primary component in striated muscle accounting for over 90% of muscle mass and volume [19]. Its integrity and homeostasis are essential for maintaining muscle function and structure after muscle injury and disease [20, 21]. MyBP-C is a key sarcomere regulatory protein, and it has been well-known for regulating muscle contraction and relaxation as well as calcium transient [1, 22]. fMyBP-C is specifically expressed in fast-twitch muscle fibers (type IIx and type IIb) which are more likely injured by mechanical stress and also challenged in diseased conditions such as muscular dystrophy and aging [4, 23, 24]. Previous studies have shown that fMyBP-C is required for peak force generation as well as muscle regeneration after injury in fast-twitch muscles [4]. However, there is still a lack of information on the expression profile of fMyBP-C in injured and diseased muscles.

The present study elucidates the crucial role of fMyBP-C in maintaining the structural integrity and functional capacity of fast-twitch muscle fibers, specifically in male skeletal muscle. The observed reductions in fMyBP-C expression in both diabetic and MDX models, as well as following eccentric contraction-induced (ECC) muscle injury, suggest that decreased fMyBP-C levels could contribute to the impaired muscle function characteristic of these conditions [25, 26]. Instead, we found elevated levels of fMyBP-C protein in the bloodstream after muscle injury, which emphasized the susceptibility of fMyBP-C to be released into circulation and suggests its possible new role as a biomarker for muscle injury such as cMyBP-C after cardiac injury [27]. The relative increase in sMyBP-C in diseased models indicates a possible compensatory mechanism; however, this adaptation appears insufficient to fully maintain muscle contractility, particularly under high-force demands. Notably, the phosphorylation profile of sMyBP-C varies significantly in response to pathophysiological stressors [28].

Our findings also highlight sex-specific variations in fMyBP-C and sMyBP-C expression and phosphorylation. Results from human muscle biopsies reveal significant differences in muscle fiber types between males and females, contributing to differences in muscle power and endurance capacity [29]. Male plantaris muscle exhibited notably higher fMyBP-C levels, aligning with prior studies that underscore the importance of this protein in rapid, high-force muscle contractions typically associated with fast-twitch muscle [4, 30]. Conversely, female muscles demonstrated significantly higher sMyBP-C phosphorylation at multiple PKA-dependent sites, suggesting that sMyBP-C phosphorylation may play a more prominent role in modulating contractility and relaxation in muscle fibers. This sex-specific expression profile also suggests that fMyBP-C may be especially critical for supporting muscle force and structural stability in male muscles, particularly under high-load conditions.

In C2^-/-^ male mice, the reduction in isometric force generation during tetanic contractions and the delayed relaxation rate imply that fMyBP-C is indispensable for optimal contractile response and recovery. Interestingly, PKA-dependent phosphorylation of sMyBP-C differed between WT and C2^-/-^ muscles, with WT samples displaying increased phosphorylation at Ser59 and Ser62 after PKA treatment, whereas C2^-/-^ samples exhibited phosphorylation shifts primarily at Ser204. This altered phosphorylation pattern in the absence of fMyBP-C suggests that the typical regulatory role of sMyBP-C may be disrupted, contributing to the observed deficits in contractile function.

Histopathological analyses of C2^-/-^ muscle fibers reveal significant adaptations, including fiber type switching from fast (type IIb) to slower fiber types (type I and IIa), accompanied by reduced cross-sectional area in type IIb fibers. Such shifts underscore the potential involvement of fMyBP-C in sustaining both the number and size of type IIb fibers, which are vital for rapid, forceful contractions. Muscle wasting or atrophy as a result of disease or aging is frequently associated with fiber-type switching and the ablation of fMyBP-C causes this phenomenon in C2^-/-^ muscles [31]. Electron microscopy further indicated structural irregularities in C2^-/-^ muscle, including expanded inter-fiber spaces and vesicle accumulation, potentially reflective of atrophic changes or chronic muscle stress [32, 33].

Transcriptomic analyses revealed substantial alterations in gene expression within C2^-/-^ muscle, with notable upregulation of *Actc1*, expressing in injured and diseased muscles, and *Phlda3*, which promotes apoptosis [34–37]. Downregulated genes included *Tm87a* and *Atp13a2*, associated with ion transport, and *Apcdd1*, a Wnt signaling inhibitor. The substantial transcriptional and functional changes collectively reinforce fMyBP-C’s essential role in regulating both the structural and contractile integrity of fast-twitch muscle fibers in male skeletal muscle.

We also highlight the essential role of fMyBP-C in sustaining muscle integrity and function as the animals age. A previous study has shown that in fast-twitch muscles the phosphorylation of sMyBP-C did not differ between adult and old mice [38]. Hence, the phenotype observed in the aging muscle cannot be attributed to sMyBP-C phosphorylation. In aged C2^-/-^ mice, the absence of fMyBP-C led to significant deficits in muscle strength and endurance, accompanied by changes in muscle fiber size and molecular signaling pathways critical for muscle homeostasis compared to WT mice. Although gross body mass and muscle weight remained unchanged, C2^-/-Old^ mice demonstrated a lower survival rate, reduced grip strength, and diminished peak plantar flexor force, indicating that fMyBP-C is vital for maintaining contractile performance in aged muscle.

The decreased force generation capacity observed in ex vivo experiments at high-frequency stimulation in C2^-/-Old^ mice muscles further supports the hypothesis that fMyBP-C is indispensable for efficient muscle contraction and force output, especially in fast-twitch fibers. These impairments in contractility were compounded by a reduced fatigue resistance, as evidenced by more rapid Po loss during repeated tetanic contractions. This reduction in endurance aligns with the established notion that age-related muscle atrophy predominantly affects fast-twitch, glycolytic fibers (type IIb), which are more dependent on fMyBP-C for rapid and forceful contractions [39].

The histological analysis provided additional insights into the structural adaptations associated with the absence of fMyBP-C in aged muscle. Although central nuclei count, an indicator of muscle regeneration did not differ between the groups, C2^-/-Old^ muscle exhibited a higher frequency of smaller muscle fibers, as indicated by the shift in fiber size distribution. This reduction in fiber size, particularly within fast-twitch types IIa, IIx, and IIb, likely contributes to the overall decline in muscle strength and highlights the importance of fMyBP-C in preserving fast-twitch fiber homeostasis during aging. The reduction in fast-twitch fiber size in C2^-/-Old^ mice further underscores the susceptibility of these fibers to atrophy in the absence of fMyBP-C, corroborating findings from prior studies on aging muscle that report selective loss of glycolytic fibers with aging [18]. Interestingly, consistent with previous literature, aging reduced the percentage of type IIb fibers in wild-type (WT) mice (young: 71.5% vs. old: 64.8%), but it did not further decrease the type IIb fiber distribution in C2KO mice (young: 58.4% vs. old: 62.4%). This may be due to an increased number of small-sized fibers in C2KO mice with aging (Figure 7C and F).

At the molecular level, our RNA Seq analysis of the fast-twitch muscle in young and aged C2^-/-^ samples showed widespread transcriptional changes (Figure S5A). Functional enrichment of commonly dysregulated genes indicates the deficiency of fMyBP-C significantly altered the skeletal muscle purine ribonucleoside catabolic process, membrane organization, actomyosin structure organization, and establishment of vesicle localization in both young and old fast-twitch fibers (supplemental file, SuppInfo.xls). Interestingly, a long noncoding RNA, B230312C02Rik is upregulated in both young and old C2^-/-^. While a noncoding RNA (2310015D24Rik) was downregulated in young mice and elevated in older mice. Based on the Mouse Genome Informatics database its expression is absent in the embryonic skeletal system and is specific to the adult skeletal system. Among the 5 genes that were up-regulated in younger but down-regulated in older C2^-/-^ mice includes two crucial genes Id-1 and Mettl21e both of which are known to have significance in cellular senescence [40], muscle fiber size and development [41, 42].

Additional enrichment analysis of age-specific DEGs in C2^-/-^ indicates that younger mice display altered molecular pathways related to apoptosis, cellular differentiation, and cellular response to oxygen levels (supplemental file, SuppInfo.xls). Disrupted pathways related to metabolic processes, membrane receptor binding, and nucleic acid regulation have been previously implicated in skeletal muscle atrophy and aging [43–45] which further emphasizes the molecular consequences of fMyBP-C absence on skeletal muscle homeostasis. As expected, similar to previous observations on skeletal gene expression transformation during aging [7, 46], pathways related to extracellular matrix organization and collagen formation were significantly altered in the older C2^-/-^ mice. These are linked to tissue repair and fibrosis and were upregulated, suggesting that the aged muscle without fMyBP-C undergoes structural remodeling likely associated with chronic stress or muscle damage. Conversely, key pathways for immune response, protein synthesis, and contractile function, including PERK signaling and skeletal muscle contraction pathways, were down-regulated, indicating a compromised ability for muscle maintenance and repair. These molecular disruptions align with the physical impairments observed and suggest that fMyBP-C may play a role in stabilizing molecular processes essential for sustaining muscle fiber composition, size, and function in aged skeletal muscle.

Interestingly, sMyBP-C protein expression increased significantly in aged C2^-/-^ muscles, consistent with observations in young C2^-/-^ mice, and may represent an adaptive response attempting to compensate for the loss of fMyBP-C. However, this upregulation does not seem sufficient to fully restore fast-twitch fiber function, indicating that the unique roles of fMyBP-C in regulating cross-bridge kinetics and sarcomere integrity are not replaceable by sMyBP-C. Moreover, an increased expression of Myomesin-1 (Myom1), a protein stabilizing the M-band of sarcomeres, was observed, potentially as a compensatory attempt to maintain structural stability in the absence of fMyBP-C.

### Conclusion

Our findings collectively provide novel insights into the sex-specific and molecular determinants of muscle functionality and highlight the critical role of fMyBP-C in preserving muscle contractility and structure, especially under high-force conditions and disease. Future studies may explore the therapeutic potential of modulating fMyBP-C expression or function in muscle disorders characterized by compromised fast-twitch fiber performance.

In conclusion, our results demonstrate that fMyBP-C is indispensable for preserving the contractile function, structural integrity, and molecular homeostasis of aged fast-twitch muscle fibers. fMyBP-C was observed to be significantly reduced in the diseased fast-twitch muscle. The absence of fMyBP-C accelerates age-related declines in muscle force production, fiber size, and regenerative capacity, emphasizing its potential as a therapeutic target to mitigate muscle aging and sarcopenia. Future research should explore interventions to maintain or enhance fMyBP-C expression or function, as this may prove beneficial for preserving muscle health, strength, and independence in the aging population.

## Supporting information

SuppInfo

## Notes

### Competing Interest Statement

The authors have declared no competing interest.

## Reference

1. Lin BL, Li A, Mun JY, Previs MJ, Previs SB, Campbell SG, et al. Skeletal myosin binding protein-C isoforms regulate thin filament activity in a Ca(2+)-dependent manner. Sci Rep. 2018;8:2604. doi:10.1038/s41598-018-21053-1

2. Giles J, Patel JR, Miller A, Iverson E, Fitzsimons D, Moss RL. Recovery of left ventricular function following in vivo reexpression of cardiac myosin binding protein C. J Gen Physiol. 2019;151:77–89. doi:10.1085/jgp.201812238

3. Lin BL, Song T, Sadayappan S. Myofilaments: Movers and Rulers of the Sarcomere. Compr Physiol. 2017;7:675–92. doi:10.1002/cphy.c160026

4. Song T, McNamara JW, Ma W, Landim-Vieira M, Lee KH, Martin LA, et al. Fast skeletal myosin-binding protein-C regulates fast skeletal muscle contraction. Proc Natl Acad Sci U S A. 2021;118:doi:10.1073/pnas.2003596118

5. Li A, Nelson SR, Rahmanseresht S, Braet F, Cornachione AS, Previs SB, et al. Skeletal MyBP-C isoforms tune the molecular contractility of divergent skeletal muscle systems. Proc Natl Acad Sci U S A. 2019;116:21882–92. doi:10.1073/pnas.1910549116

6. Li M, Andersson-Lendahl M, Sejersen T, Arner A. Knockdown of fast skeletal myosin-binding protein C in zebrafish results in a severe skeletal myopathy. J Gen Physiol. 2016;147:309–22. doi:10.1085/jgp.201511452

7. Lin IH, Chang JL, Hua K, Huang WC, Hsu MT, Chen YF. Skeletal muscle in aged mice reveals extensive transformation of muscle gene expression. BMC Genet. 2018;19:55. doi:10.1186/s12863-018-0660-5

8. Ochala J, Frontera WR, Dorer DJ, Van Hoecke J, Krivickas LS. Single skeletal muscle fiber elastic and contractile characteristics in young and older men. J Gerontol A Biol Sci Med Sci. 2007;62:375–81. doi:10.1093/gerona/62.4.375

9. Song T, Manoharan P, Millay DP, Koch SE, Rubinstein J, Heiny JA, et al. Dilated cardiomyopathy-mediated heart failure induces a unique skeletal muscle myopathy with inflammation. Skelet Muscle. 2019;9:4. doi:10.1186/s13395-019-0189-y

10. Brooks SV, Faulkner JA. Contractile properties of skeletal muscles from young, adult and aged mice. J Physiol. 1988;404:71–82. doi:10.1113/jphysiol.1988.sp017279

11. Zhou Y, Zhou B, Pache L, Chang M, Khodabakhshi AH, Tanaseichuk O, et al. Metascape provides a biologist-oriented resource for the analysis of systems-level datasets. Nat Commun. 2019;10:1523. doi:10.1038/s41467-019-09234-6

12. Haizlip KM, Harrison BC, Leinwand LA. Sex-based differences in skeletal muscle kinetics and fiber-type composition. Physiology (Bethesda). 2015;30:30–9. doi:10.1152/physiol.00024.2014

13. Colson BA, Locher MR, Bekyarova T, Patel JR, Fitzsimons DP, Irving TC, et al. Differential roles of regulatory light chain and myosin binding protein-C phosphorylations in the modulation of cardiac force development. J Physiol. 2010;588:981–93. doi:10.1113/jphysiol.2009.183897

14. Ackermann MA, Kontrogianni-Konstantopoulos A. Myosin binding protein-C slow is a novel substrate for protein kinase A (PKA) and C (PKC) in skeletal muscle. J Proteome Res. 2011;10:4547–55. doi:10.1021/pr200355w

15. Trappe S, Williamson D, Godard M. Maintenance of whole muscle strength and size following resistance training in older men. J Gerontol A Biol Sci Med Sci. 2002;57:B138–43. doi:10.1093/gerona/57.4.b138

16. Tieland M, Trouwborst I, Clark BC. Skeletal muscle performance and ageing. J Cachexia Sarcopenia Muscle. 2018;9:3-19. doi:10.1002/jcsm.12238

17. Larsson L, Degens H, Li M, Salviati L, Lee YI, Thompson W, et al. Sarcopenia: Aging-Related Loss of Muscle Mass and Function. Physiol Rev. 2019;99:427–511. doi:10.1152/physrev.00061.2017

18. Seo JY, Kang JS, Kim YL, Jo YW, Kim JH, Hann SH, et al. Maintenance of type 2 glycolytic myofibers with age by Mib1-Actn3 axis. Nat Commun. 2021;12:1294. doi:10.1038/s41467-021-21621-6

19. Swist S, Unger A, Li Y, Voge A, von Frieling-Salewsky M, Skarlen A, et al. Maintenance of sarcomeric integrity in adult muscle cells crucially depends on Z-disc anchored titin. Nat Commun. 2020;11:4479. doi:10.1038/s41467-020-18131-2

20. Martin TG, Kirk JA. Under construction: The dynamic assembly, maintenance, and degradation of the cardiac sarcomere. J Mol Cell Cardiol. 2020;148:89–102. doi:10.1016/j.yjmcc.2020.08.018

21. Proske U, Morgan DL. Muscle damage from eccentric exercise: mechanism, mechanical signs, adaptation and clinical applications. J Physiol. 2001;537:333–45. doi:10.1111/j.1469-7793.2001.00333.x

22. Moss RL, Fitzsimons DP, Ralphe JC. Cardiac MyBP-C regulates the rate and force of contraction in mammalian myocardium. Circ Res. 2015;116:183–92. doi:10.1161/CIRCRESAHA.116.300561

23. Gehrig SM, Koopman R, Naim T, Tjoakarfa C, Lynch GS. Making fast-twitch dystrophic muscles bigger protects them from contraction injury and attenuates the dystrophic pathology. Am J Pathol. 2010;176:29–33. doi:10.2353/ajpath.2010.090760

24. Ferrucci L, Baroni M, Ranchelli A, Lauretani F, Maggio M, Mecocci P, et al. Interaction between bone and muscle in older persons with mobility limitations. Curr Pharm Des. 2014;20:3178–97. doi:10.2174/13816128113196660690

25. Hammers DW, Hart CC, Matheny MK, Wright LA, Armellini M, Barton ER, et al. The D2.mdx mouse as a preclinical model of the skeletal muscle pathology associated with Duchenne muscular dystrophy. Sci Rep. 2020;10:14070. doi:10.1038/s41598-020-70987-y

26. Espino-Gonzalez E, Dalbram E, Mounier R, Gondin J, Farup J, Jessen N, et al. Impaired skeletal muscle regeneration in diabetes: From cellular and molecular mechanisms to novel treatments. Cell Metab. 2024;36:1204–36. doi:10.1016/j.cmet.2024.02.014

27. Kuster DW, Cardenas-Ospina A, Miller L, Liebetrau C, Troidl C, Nef HM, et al. Release kinetics of circulating cardiac myosin binding protein-C following cardiac injury. Am J Physiol Heart Circ Physiol. 2014;306:H547–56. doi:10.1152/ajpheart.00846.2013

28. Ackermann MA, Kerr JP, King B, C WW, Kontrogianni-Konstantopoulos A. The Phosphorylation Profile of Myosin Binding Protein-C Slow is Dynamically Regulated in Slow-Twitch Muscles in Health and Disease. Sci Rep. 2015;5:12637. doi:10.1038/srep12637

29. Nuzzo JL. Sex differences in skeletal muscle fiber types: A meta-analysis. Clin Anat. 2024;37:81–91. doi:10.1002/ca.24091

30. Song T, Landim-Vieira M, Ozdemir M, Gott C, Kanisicak O, Pinto JR, et al. Etiology of genetic muscle disorders induced by mutations in fast and slow skeletal MyBP-C paralogs. Exp Mol Med. 2023;55:502–9. doi:10.1038/s12276-023-00953-x

31. Ciciliot S, Rossi AC, Dyar KA, Blaauw B, Schiaffino S. Muscle type and fiber type specificity in muscle wasting. Int J Biochem Cell Biol. 2013;45:2191–9. doi:10.1016/j.biocel.2013.05.016

32. Wang W, Li M, Chen Z, Xu L, Chang M, Wang K, et al. Biogenesis and function of extracellular vesicles in pathophysiological processes of skeletal muscle atrophy. Biochem Pharmacol. 2022;198:114954. doi:10.1016/j.bcp.2022.114954

33. Bonaldo P, Sandri M. Cellular and molecular mechanisms of muscle atrophy. Dis Model Mech. 2013;6:25–39. doi:10.1242/dmm.010389

34. Hicks MR, Saleh KK, Clock B, Gibbs DE, Yang M, Younesi S, et al. Regenerating human skeletal muscle forms an emerging niche in vivo to support PAX7 cells. Nat Cell Biol. 2023;25:1758–73. doi:10.1038/s41556-023-01271-0

35. Chong JX, Childers MC, Marvin CT, Marcello AJ, Gonorazky H, Hazrati LN, et al. Variants in ACTC1 underlie distal arthrogryposis accompanied by congenital heart defects. HGG Adv. 2023;4:100213. doi:10.1016/j.xhgg.2023.100213

36. Sztal TE, McKaige EA, Williams C, Ruparelia AA, Bryson-Richardson RJ. Genetic compensation triggered by actin mutation prevents the muscle damage caused by loss of actin protein. PLoS Genet. 2018;14:e1007212. doi:10.1371/journal.pgen.1007212

37. Kim S, Ayan B, Shayan M, Rando TA, Huang NF. Skeletal muscle-on-a-chip in microgravity as a platform for regeneration modeling and drug screening. Stem Cell Reports. 2024;19:1061–73. doi:10.1016/j.stemcr.2024.06.010

38. Perazza LR, Wei G, Thompson LV. Fast and slow skeletal myosin binding protein-C and aging. Geroscience. 2023;45:915–29. doi:10.1007/s11357-022-00689-y

39. Akasaki Y, Ouchi N, Izumiya Y, Bernardo BL, Lebrasseur NK, Walsh K. Glycolytic fast-twitch muscle fiber restoration counters adverse age-related changes in body composition and metabolism. Aging Cell. 2014;13:80–91. doi:10.1111/acel.12153

40. Alani RM, Young AZ, Shifflett CB. Id1 regulation of cellular senescence through transcriptional repression of p16/Ink4a. Proc Natl Acad Sci U S A. 2001;98:7812–6. doi:10.1073/pnas.141235398

41. Wang C, Zhang B, Ratliff AC, Arrington J, Chen J, Xiong Y, et al. Methyltransferase-like 21e inhibits 26S proteasome activity to facilitate hypertrophy of type IIb myofibers. FASEB J. 2019;33:9672–84. doi:10.1096/fj.201900582R

42. Gundersen K, Merlie JP. Id-1 as a possible transcriptional mediator of muscle disuse atrophy. Proc Natl Acad Sci U S A. 1994;91:3647–51. doi:10.1073/pnas.91.9.3647

43. Mielcarek M, Smolenski RT, Isalan M. Transcriptional Signature of an Altered Purine Metabolism in the Skeletal Muscle of a Huntington’s Disease Mouse Model. Front Physiol. 2017;8:127. doi:10.3389/fphys.2017.00127

44. Zoref-Shani E, Shainberg A, Sperling O. Alterations in purine nucleotide metabolism during muscle differentiation in vitro. Biochem Biophys Res Commun. 1983;116:507–12. doi:10.1016/0006-291x(83)90552-1

45. Miller SG, Hafen PS, Brault JJ. Increased Adenine Nucleotide Degradation in Skeletal Muscle Atrophy. Int J Mol Sci. 2019;21:doi:10.3390/ijms21010088

46. Lofaro FD, Cisterna B, Lacavalla MA, Boschi F, Malatesta M, Quaglino D, et al. Age-Related Changes in the Matrisome of the Mouse Skeletal Muscle. Int J Mol Sci. 2021;22:doi:10.3390/ijms221910564

