## Supplemental Figures for "Fast Myosin Binding Protein-C is a Vital Regulator in Young and Aged Fast Skeletal Muscle Homeostasis"

Akhil Baby et al.,

The PDF file includes: Figures S1 to S8

Figure S1

A

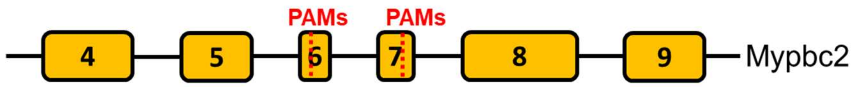


Exon6:
CACCCCGTCAGGACTCAT**CCG**GTCAGAGCCTCGAGAGCTTCAAGCGTTC
Exon7:
GGGTGACGGGAAGTCAGAAGATGCAGGCGAGCTGGATTTCAG**TGG**CTTGTTGAAGAAGAG

B C

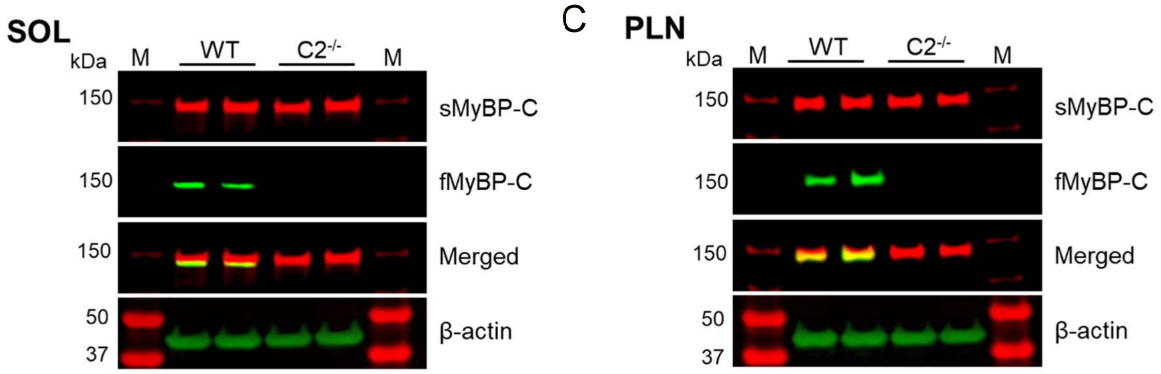


**Figure S1. Second *Mybpc2* knockout mouse model generation.** A. Wildtype Mybpc2 sequence targeting exon 6 and 7 using CRISPR/Cas9 system. PAM sites for sgRNAs are highlighted in red. B. Complete knockdown of fMyBP-C protein was confirmed in both slow (soleus, SOL) and fast (plantaris, PLN) twitch muscles of the *Mybpc2* knockout mice by western blot analysis. n = 2 muscle samples.

### Figure S2

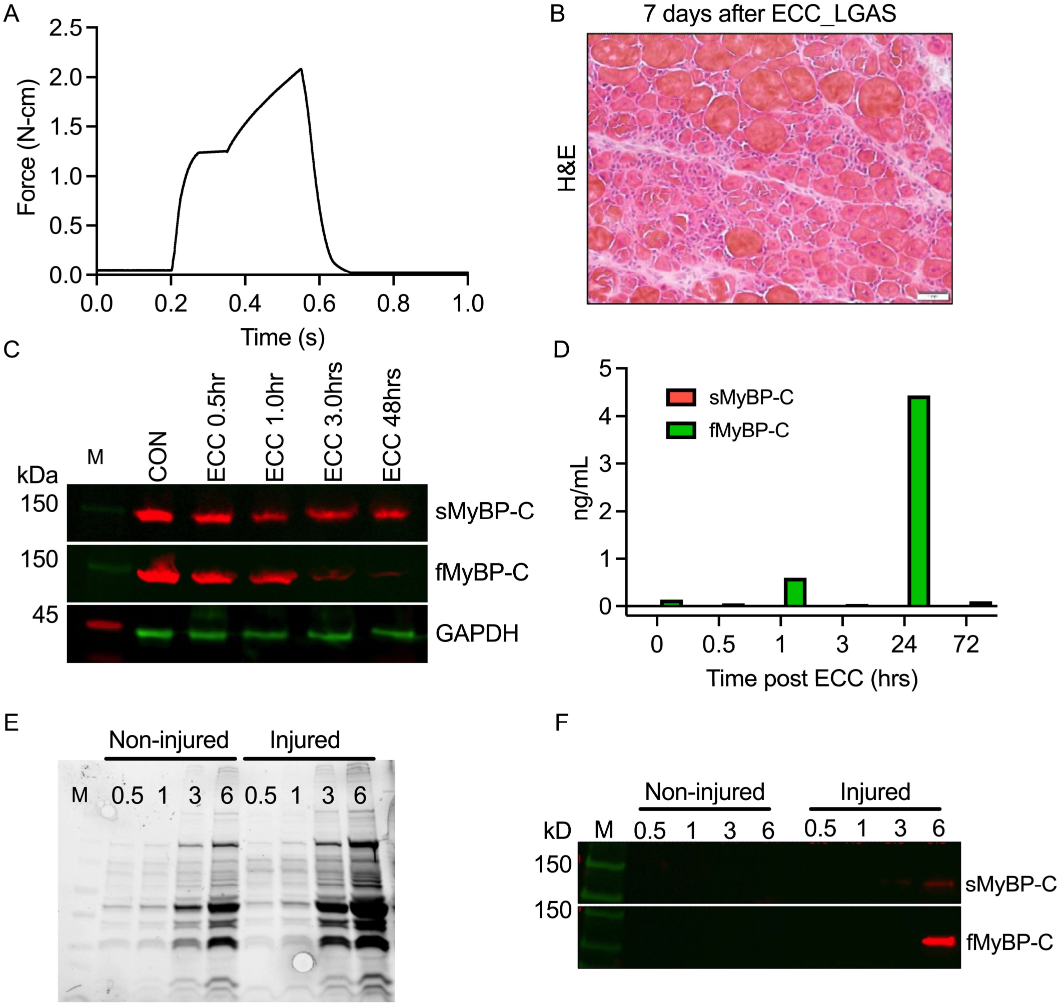


**Figure S2. Loss of fMyBP-C after muscle injury.** (A) Force-time graph during eccentric muscle contraction (ECC) of plantar flexor muscle. (B) Cross-sectioned lateral gastrocnemius muscle (LGAS) stained with H&E at 7 days after ECC injury. (C) Decreased fMyBP-C expression post-ECC induced muscle injury. (D) ELISA assay detected elevated fMyBP-C levels in the blood after ECC injury. One day after ECC contraction, the GAS muscle was dissected and incubated in 800uL PBS solution. 100uL of effluent was collected at 0.5, 1.0, 3.0, and 6.0 hours after incubation. (E) Coomassie-stained gel image loaded with 10uL effluent. (F) Slow and fast MyBP-C were detected in the ECC injured effluent incubated for 6.0 hours.

#### Figure S3

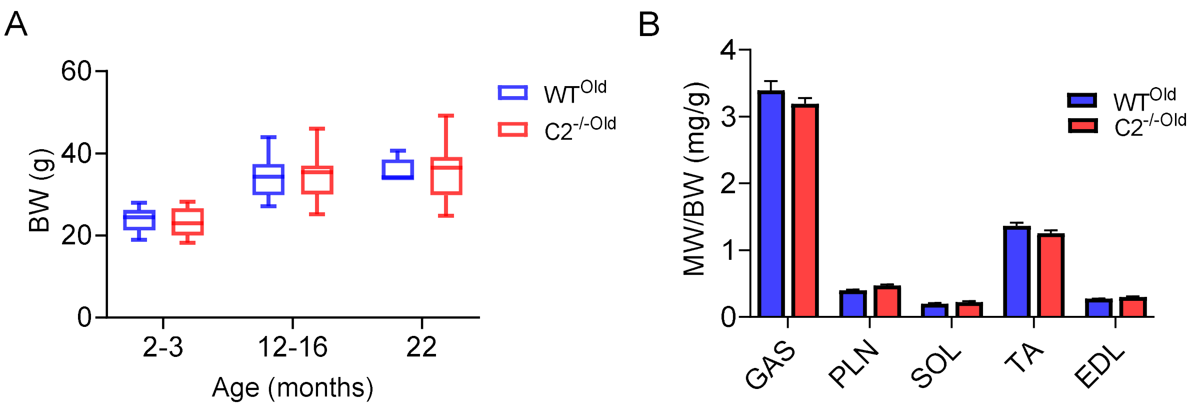


**Figure S3. Preserved body and muscle weight in C2$^{-/-Old}$ mice.** Absolute body weight (A) and normalized hindlimb muscle mass (B) by body weight at 22 months were not significantly different between WT (n = 6~16) and C2$^{-/-}$ (n = 15~16) mice. Error bars represent ± SEM.

### Figure S4

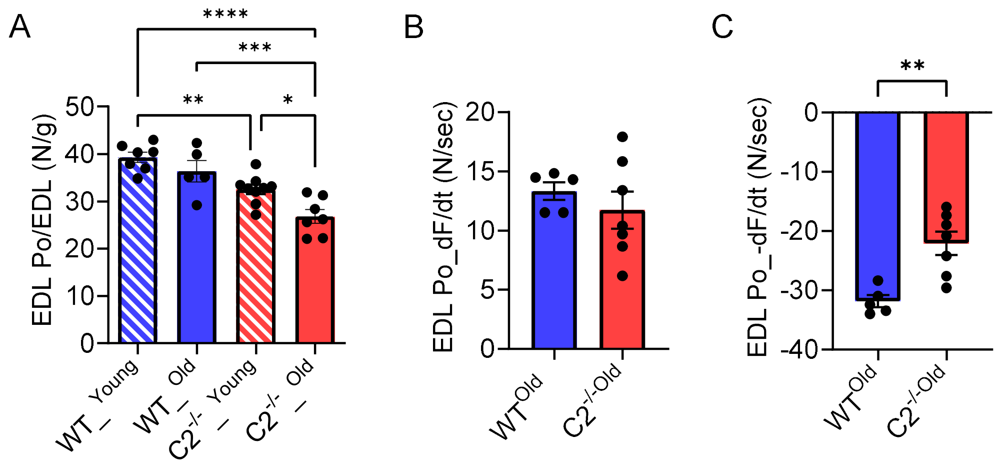


**Figure S4. Reduced $C2^{-/-}$ EDL muscle functions with aging.** (A) Peak isometric tetanic force (Po) of young and old WT and $C2^{-/-}$ EDL muscles. Rate of activation (B) and relaxation (C) during $P_o$ generation in aged WT and $C2^{-/-}$. n = 5-9 muscles, **P*<0.05, ***P*<0.01, ****P*<0.001, and *****P*<0.0001, comparing each group to every other group.

### Figure S5

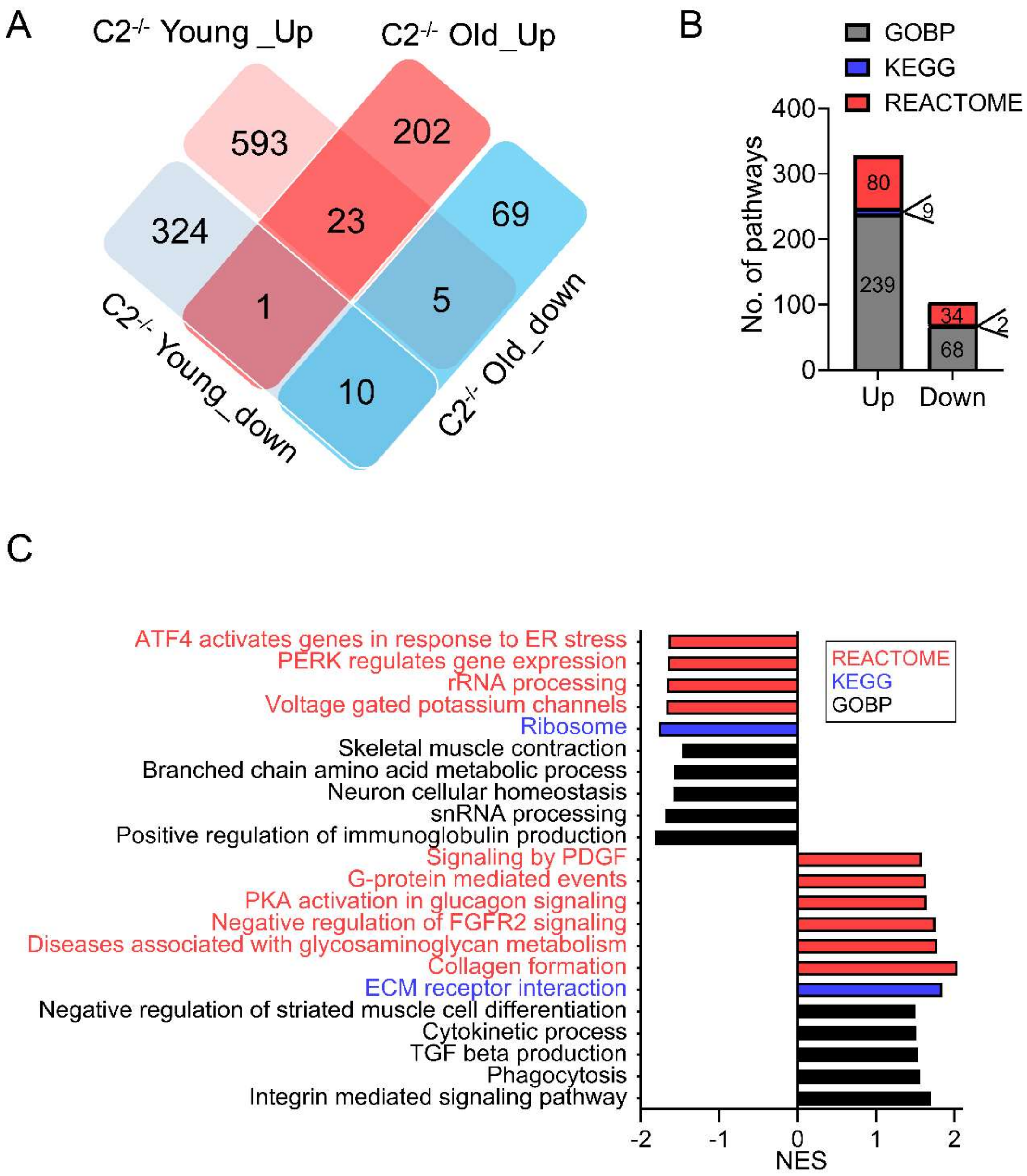


**Figure S5. Comparative analysis of young and old fast twitch muscle transcriptome in absence of fMyBP-C.** The RNA sequencing data from the young and old fast twitch muscle fibers were compared to identify the C2$^{-/-}$ and age specific alterations of muscle transcriptome. (A) Venn diagram displaying the number of genes that display C2$^{-/-}$ and age specific dysregulation in C2$^{-/-}$ muscle. (B) Total upregulated and downregulated signals of C2$^{-/-}$ DEGs in old mice analyzed by gene set enrichment assay (GSEA) with GOBP, KEGG, and REACTOME database. (C) Selected most upregulated and downregulated pathways in C2$^{-/-}$ old mice.

### Figure S6

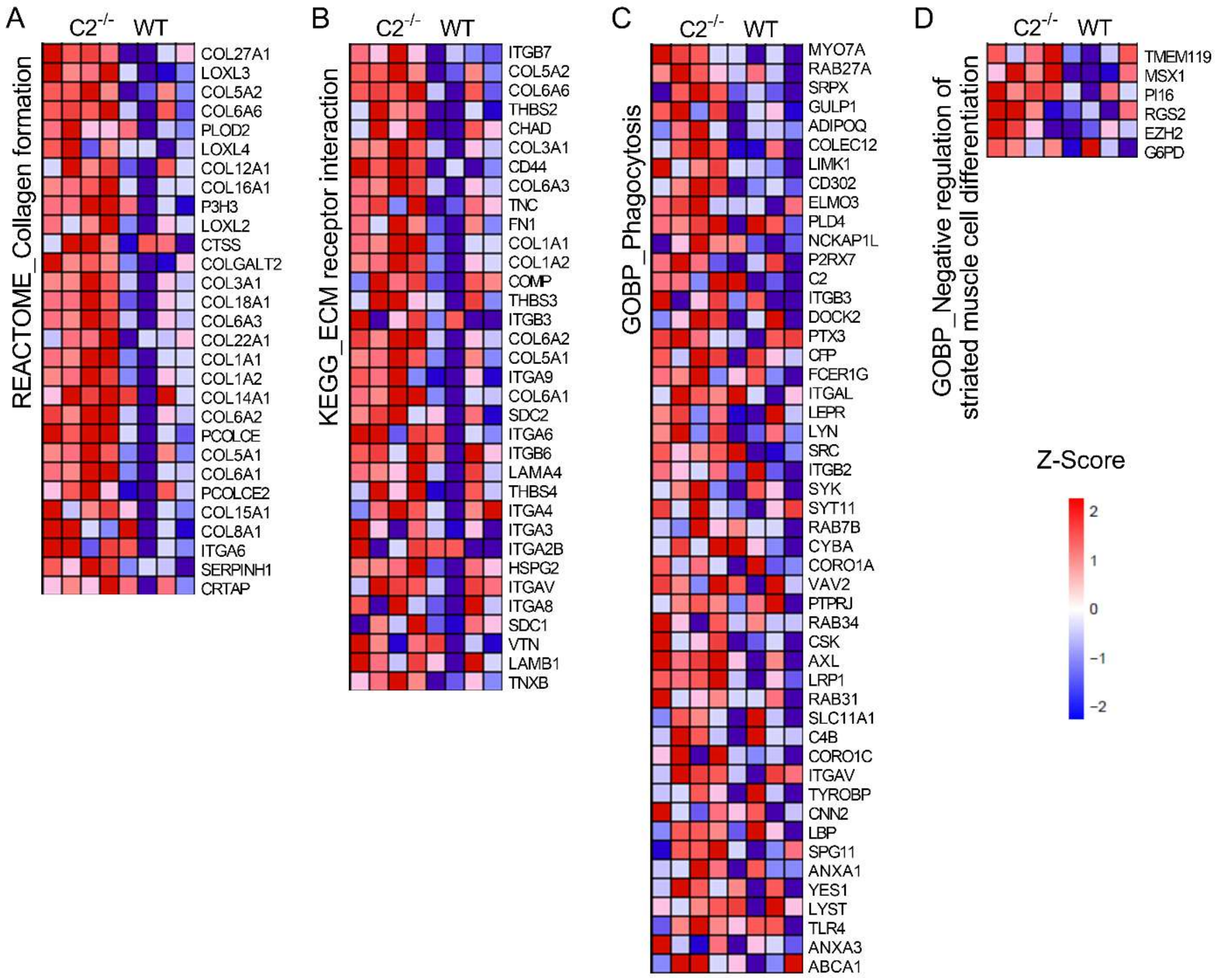


**Figure S6. Heat map of upregulated core enrichment genes in aged $C2^{-/-}$ selected by GSEA.** (A) REACTOME_Collagen formation. (B) KEGC_ECM receptor interaction. (C) GOBP_Phagocytosis. (D) GOBP_Negative regulation of striated muscle cell differentiation. n = 4 TA samples.

### Figure S7

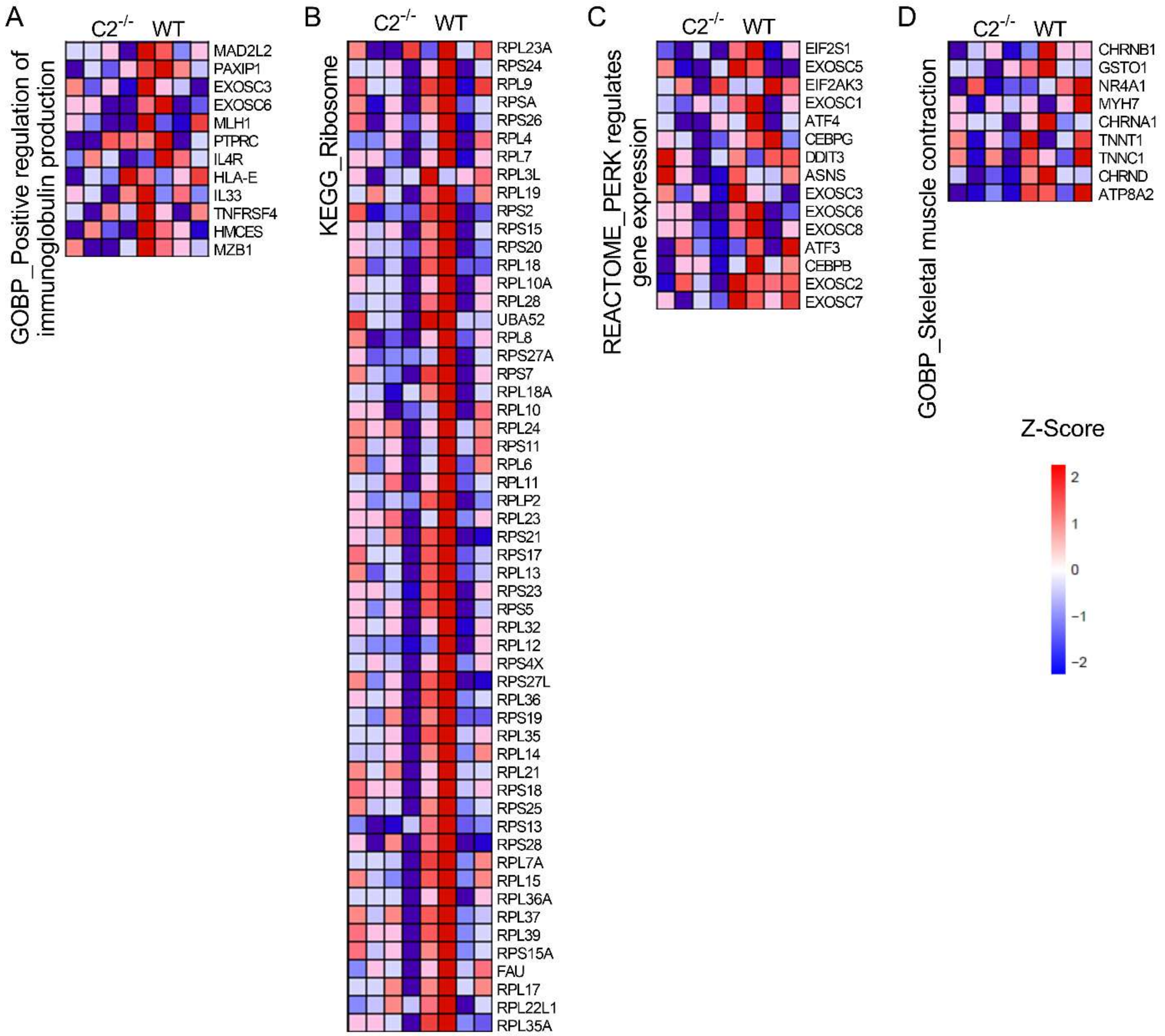


**Figure S7. Heat map of downregulated core enrichment genes in aged C2$^{-/-}$ selected by GSEA.** (A) GOBP_Positive regulation of immunoglobulin production. (B) KEGG_Ribosome. (C) REACTOME_PERK regulates gene expression. (D) GOBP_Skeletal muscle contraction. n=4 TA samples.

### Figure S8

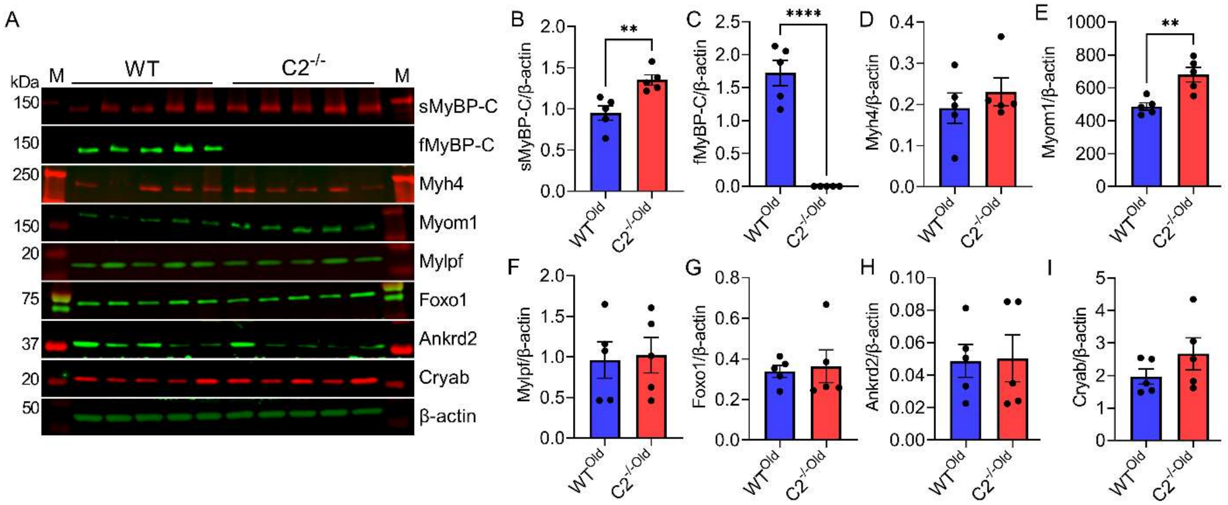


**Figure S8. Expression of key sarcomere structure and muscle atrophy-related proteins in aged WT and C2$^{-/-}$ TA muscle.** Western blot images (A) and quantification of sarcomere protein (sMyBP-C (B), fMyBP-C (C), Myh4 (D), Myom1 (E), Mylpf (F) and atrophy-related proteins (Foxo1 (G), Ankrd2 (H), Cryab (I)) expressions. Error bars mean ± SEM and ***P*<0.01 and *****P*<0.0001 to C2$^{-/-Old}$ vs. WT$^{Old}$. n = 5 mice muscle samples.
